# Positional ^13^C Enrichment Analysis of Aspartate by GC-MS to Determine PEPC Activity *In Vivo*

**DOI:** 10.1101/2024.05.07.592938

**Authors:** Luisa Wittemeier, Yogeswari Rajarathinam, Alexander Erban, Martin Hagemann, Joachim Kopka

**Affiliations:** Max-Planck Institute of Molecular Plant Physiology, Am Mühlenberg 1, 14476 Potsdam, Germany; Department of Plant Physiology, University of Rostock, Albert-Einstein-Straße 3, 18059 Rostock, Germany

**Author notes:** **Corresponding Authors** Luisa Wittemeier - Max-Planck Institute of Molecular Plant Physiology, Am Mühlenberg 1, 14476 Potsdam, Germany;, Joachim Kopka - Max-Planck Institute of Molecular Plant Physiology, Am Mühlenberg 1, 14476 Potsdam, Germany. Yogeswari Rajarathinam - Environmental and Biochemical Sciences, James Hutton Institute, Invergowrie, Dundee, DD2 5DA, Scotland, UK.

## Abstract

Photoautotrophic organisms fix inorganic carbon (Ci) by two enzymes, ribulose-1,5-bisphosphate carboxylase/oxygenase (RUBISCO) and phosphoenolpyruvate carboxylase (PEPC). RUBISCO assimilates Ci (CO2) into the 1-C position of 3-phosphoglycerate (3PGA). The Calvin-Benson-Basham (CBB) cycle redistributes fixed carbon atoms into 2,3-C_2_ of the same molecule. PEPC uses phosphoenolpyruvate (PEP) derived from 3PGA and assimilates Ci (HCO_3-_) into 4-C of oxaloacetate (OAA). 1,2,3-C_3_ of OAA and of its transaminase product aspartate originate directly from 1,2,3-C_3_ of 3PGA. Positional isotopologue analysis of aspartate, the main downstream metabolite of OAA in the model cyanobacterium *Synechocystis* sp. PCC 6803 (*Synechocystis*), allows differentiation between PEPC, RUBISCO, and CBB cycle activities within one molecule. We explored *in source* fragmentation of gas chromatography-electron impact ionization-mass spectrometry (GC-EI-MS) at nominal mass resolution and GC-atmospheric pressure chemical ionization-MS (GC-APCI-MS) at high mass resolution. This enabled the determination of fractional ^13^C enrichment (E^13^C) at each carbon position of aspartate. Two prevailing GC-MS derivatization methods, i.e. trimethylsilylation and tert-butyldimethylsilylation, were evaluated. The method was validated by ^13^C-isotopomer mixtures of positional labeled aspartic acid. Combination with dynamic ^13^CO_2_ labeling of *Synechocystis* cultures allowed direct measurements of PEPC activity *in vivo* alongside analyses of RUBISCO and CBB cycle activities. Accurate quantification of aspartate concentration and positional E^13^C provided molar Ci assimilation rates during the day and night phases of photoautotrophic *Synechocystis* cultures. The validated method offers several applications to characterize the photosynthetic Ci fixation in different organisms.

## Introduction

Metabolic flux analyses reveal changes of metabolism. ^13^C labelling and analysis by gas chromatography mass spectrometry (GC-MS) are valuable techniques to examine production and consumption rates of metabolites. Labelling can either be analyzed at isotopic steady-state to verify contributions of different carbon sources, i.e. alternative uses of metabolic pathways and reactions, or with dynamic flux studies, in which concentration and labelling information of metabolites are monitored at consecutive timepoints after the stable isotope pulse. In photoautotrophic metabolism, CO_2_ is the only substrate which results in uniformly labeled metabolites under metabolic steady-state. By contrast, dynamic flux studies monitor the rates of label incorporation^1^ where the initial labelling can be non-uniform depending on enzyme reaction mechanisms. However, measurements of ^13^C enrichment into a complete molecule does not allow direct statements about the source of carbon assimilation into this molecule if more than one reaction contributes to its formation.

Positional ^13^C labelling information is crucial to understand metabolic pathway regulation and to estimate enzyme activities *in vivo*. Positional information after ^13^C labeling can be directly obtained by nuclear magnetic resonance spectroscopy (NMR)^2^. This versatile quantitative technology is limited by low sensitivity and requires large amounts of biological material. GC-MS is equally widely established for metabolomic analyses, offers higher sensitivity and provides information at isotopologue level. MS determines how many carbons of a molecule are labeled, but does not allow direct conclusions on positional labeling and the presence of defined isotopomers, i.e. isomers with identical isotopic composition but isotopes at different positions of the structure^3, 4^. *In silico* fragmentation analysis of positional labeled TCA cycle metabolites analyzed by GC-EI-MS and GC-EI-MS/MS suggested several mass fragments that are useful for positional fractional ^13^C enrichment (E^13^C) analysis^5^. Several previous studies showed that calculation of positional E^13^C is possible using GC-EI-MS measurements of amino acids, organic acids or glucose^6,^ _7_.

Photoautotrophic organisms such as cyanobacteria, algae, and plants use oxygenic photosynthesis to produce reducing power (NADPH) and energy (ATP) in the light reactions with oxygen as a by-product. These products from light reactions drive inorganic carbon (Ci) fixation and production of organic biomass. In cyanobacteria, Ci, in forms of CO_2_ or HCO_3_^-^, enters metabolism via two main carboxylation reactions that are catalyzed by RUBISCO and PEPC. RUBISCO, the key enzyme of the Calvin-BensonBassham (CBB) cycle, assimilates CO_2_ using ribulose-1,5-bisphosphate (RuBP) and generates two molecules 3-phosphoglycerate (3PGA). PEPC fixes HCO_3_^-^ using phosphoenolpyruvate (PEP) and forms oxaloacetate (OAA) and inorganic phosphate. OAA is immediately converted to malate, aspartate or citrate by malate dehydrogenase (MDH), aspartate aminotransferase (AAT) or citrate synthase (CS), respectively. The carbon backbone of OAA and of the downstream metabolites, aspartate or malate, comprises four carbon atoms. 1,2,3-C_3_ is derived from PEP that maintains the carbon constellation of 3PGA from Ci assimilation via RUBISCO and the CBB cycle. 4-C is added by PEPC^8^.

In land plants, PEPC is crucial for C_4_ and CAM photosynthesis^9^. Malate or aspartate serve as transport metabolites in variants of C_4_ metabolism. CAM photosynthesis uses malate to build a temporal carbon store in vacuoles during the night that delivers CO_2_ to RUBISCO by the action of malic enzyme during the day. Through these mechanisms, Ci is concentrated in the vicinity of RUBISCO and the wasteful RUBISCO oxygenase reaction suppressed. PEPC has an alternative key function by catalyzing the anaplerotic synthesis of OAA to replenish TCA cycle intermediates^10^. It has been reported that PEPC greatly contributes to Ci assimilation in addition to RUBISCO in cyanobacteria^11^. For example, PEPC has a substantial contribution to CO_2_ assimilation yielding 25% of total Ci assimilation in *Synechocystis* sp. PCC 6803 (*Synechocystis*) under mixotrophic conditions as was suggested by metabolic flux studies based on ^13^C-glucose labelling^12^ or 12% under photoautotrophic conditions as was predicted by genome-scale isotopic non-stationary metabolic flux analysis^13^. PEPC is an essential enzyme in *Synechococcus* PCC 7942 and *Synechocystis*^14, 15^. PEPC orthologs of more evolved cyanobacterial clades, such as *Oscillatoriales* and *Nostocales*, have characteristic features that are comparable to C_4_ isoforms, whereas more basal groups of cyanobacteria, such as *Chroococcales* and *Pleurocapsales*, have neither dedicated C_3_ nor C_4_ isoforms^16^.

PEPC activity can be determined *in vitro* by spectrophotometric assays that are coupled to MDH and lactate dehydrogenase and are based on measurements of NADH reduction^17^. *In vitro* activity assays do not account for cellular cofactors, regulators, substrate availability, effects of subcellular structures or metabolic channeling. Hence, those tests will never reflect cytosolic *in vivo* conditions completely. To address this analytical gap, we have undertaken to assess PEPC activity *in vivo* by mass spectrometry. Direct analysis of OAA is demanding due to its very low cellular concentration and its rapid conversion into downstream metabolites, e.g. aspartate, malate, and citrate^18, 19^. In land plants, positional isotopomer analysis of malate by either NMR or GC-MS was used as a proxy for PEPC activity *in vivo* ^20, 21^. However, flux balance analysis suggested that the main flux in *Synechocystis* leads from OAA via AAT to aspartate^22^. Non-stationary metabolic flux analyses detected flux in photoautotrophic conditions through MDH, AAT and CS, whereby MDH and AAT seem to catabolize OAA to equivalent amounts^13, 23^. The modelled ranges of fluxes through many enzyme reactions were, however, still very wide^13^. *Synechocystis* CS has low catalytic activity and seems to be an inefficient enzyme causing low flux through the oxidative branch of the tricarboxylic acid (TCA) cycle^24^.

Previously, GC and LC coupled to tandem mass spectrometry (MS/MS) allowed determination of the complete isotopomer distribution of aspartate^25, 26^. Here, we applied routine direct *in-source* fragmentation of GC-MS systems. We predicted aspartic acid fragmentation *in silico* and confirmed predictions through analysis of positional labeled aspartic acid reference substances in combination with exact mass determinations. These analyses were enabled by a GC-APCI-MS instrument. Limitations of detection and accuracy of ^13^C enrichment measurements were evaluated based on mixtures of positional labeled and natural aspartic acid. In combination with dynamic ^13^CO_2_ labelling of *Synechocystis*, we determined position specific carbon assimilation into each carbon atom of aspartate and assessed proxies of PEPC activity *in vivo* alongside contributions of RUBISCO and CBB cycle activities during day light and night conditions. Our method proves the expected absence of RUBISCO activity in the dark and changing PEPC activity upon day to night shift.

### Experimental Section Aspartic acid standard mixtures

Natural aspartic acid, fully labeled [U-^13^C]aspartic acid and positional labeled [1-^13^C], [2-^13^C], [3-^13^C], [4-^13^C]aspartic acid were purchased from Sigma Aldrich/Merck KGaA (Darmstadt, Germany), according to analytical certificates of the manufacturer with natural, 99.0, 99.6, 99.7, 99.6, and 99.3% ^13^C enrichment of the complete molecules and at the single carbon positions, respectively. Mixtures with different ratios of natural and all positional labeled aspartic acids were prepared in aqueous solution at molar ratios of 5:95, 10:90, 50:50, 90:10, or 95:5 and dried by vacuum centrifugation (Table S1). The dried mixtures were used to assess the accuracy of E^13^C determination at 25 ng per GC-MS injection as the mean deviation of measured E^13^C from expected E^13^C and the precision of E^13^C measurements as the standard deviation (SD) of E^13^C deviation. To determine the lower detection threshold of E^13^C measurements, positional labeled aspartic acid standards were mixed in a molar ratio of 1:1:1:1. This mixture was diluted by natural aspartic acid to 100, 50, 20, 10 and 4% of total aspartic acid (Table S1). Different amounts of the isotope-diluted mixtures were analyzed by GC -MS at 1.25, 2.5, 12.5, 25, 125 and 250 ng per injection. All samples were spiked with 25 ng per injection of ^13^C_6_-sorbitol (Sigma-Aldrich/Merck KGaA, 99 atom % ^13^C, 99%) as constant internal standard.

### Cultivation and sampling of *Synechocystis* sp. PCC 6803

Glucose-tolerant *Synechocystis* wild-type (WT) precultures were grown for 3 days at 30°C in modified BG11 growth medium^27^. To avoid an additional carbon source, citric acid and ferric ammonium citrate were replaced by a 0.021 mM FeCl_3_ iron source. *Synechocystis* cells were cultivated in a Multicultivator MC 1000-OD photobioreactor (Photon Systems Instruments, Drásov, Czech Republic) under continuous illumination set to 100 μmol photons m^-2^ s^-1^ and with high inorganic carbon supply, i.e. constant bubbling (2 bubbles * s^-1^) with 5% CO_2_ in air. After pre-cultivation, culture media were renewed by centrifugation and resuspension of cells. Illumination was changed to 12 h light/12 h dark cycles maintaining 100 μmol photons m^-2^ s^-1^ in the light phases. After three day/night cycles medium was exchanged and OD_750_ adjusted to ∼0.8 in the third light phase 4 h prior to the onset of darkness by fast vacuum filtration with continuous illumination. Sampling of cells was at and after transition to the fourth night by <15 s filtration onto glass fiber filters (25 mm, pore size 1.6 μm, Cytiva, Sigma Aldrich/Merck KGaA) and immediate shock freezing in liquid N_2_. First sampling at t_0_ was harvested in the light with 5% ambient CO_2_ bubbling. Prior to dynamic ^13^CO_2_ labelling at the onset of darkness, culture medium was exchanged by fast filtration to remove dissolved non-labeled Ci from the cultures. Cells were resuspended in the dark by immediate bubbling with 5% ^13^CO_2_ in synthetic air. Labeled samples at t_1_-t_6_ were collected 5, 10, 15, 30 and 90 min after beginning of ^13^CO_2_ bubbling. OD_750_ mL^-1^ was adjusted to ∼1.0 and recorded of all individual cultures at t_0_, t_1_ and t_6_. For continuous light cultivation and dynamic ^13^CO_2_ labelling in the light, the procedure was identical omitting only the change to day night cycles.

### Metabolite extraction

Polar metabolites were extracted from deep frozen cells on filters using methanol (Sigma-Aldrich, gradient grade for liquid chromatography, ≥99.9%) chloroform (Sigma-Aldrich/Merck KGaA, contains ethanol as stabilizer, ACS reagent grade, ≥99.8%) and double distilled water (ddH_2_O). 1 mL of extraction mix, i.e. methanol:chloroform:ddH_2_O (2.5:1:1; v/v/v) with 6 μg * mL^-1 13^C_6_-sorbitol as internal standard, was added and incubated 15 min at 70°C under permanent agitation. Phase separation was induced by adding 400 μL of ddH_2_O. The upper polar phase (∼800 μL) was separated by centrifugation and dried overnight by vacuum centrifugation^28^.

### Chemical derivatization for GC-MS analysis

Chemical derivatization of dried metabolite samples for GC-MS analysis was exactly as described previously omitting 4-(dimethylamino)pyridine^29^. Samples were subjected to methoxyamination followed by either trimethylsilylation (TMS) using *N,O*-bis(trimethylsilyl)trifluoroacetamide (BSTFA, MachereyNagel, Düren, Germany) or using the same protocol replacing trimethylsilylation by *tert*.-butyldimethylsilylation (TBDMS) with *N*-methyl-*N*-(*tert*.-butyldimethylsilyl)trifluoroacetamide (MBDSTFA, Macherey-Nagel) reagent and an incubation for 60 min at 70°C followed by 15 min at 30°C under permanent agitation.

### GC-MS analysis

Derivatized samples were analyzed by an Agilent 6890N24 gas chromatograph (Agilent Technologies, Waldbronn, Germany) hyphenated to either electron impact ionization-time of flight-mass spectrometry (EI-TOF-MS) using a LECO Pegasus III time of flight mass spectrometer (LECO Instrumente GmbH, Mönchengladbach, Germany) or to atmospheric pressure chemical ionization-time of flight-mass spectrometry (APCI-TOF-MS) with a micrOTOF-Q II hybrid quadrupole time-of-flight mass spectrometer (Bruker Daltonics, Bremen, Germany) equipped with an APCI ion source and GC interface (Bruker Daltonics)^30^. All measurements were conducted in splitless mode using 5% phenyl 95% dimethylpolysiloxane fused silica capillary column with 30 m length, 0.25 mm inner diameter, 0.25 μm film thickness and an integrated 10 m precolumn (Agilent Technologies (CP9013)). Measurements of chemical standard mixtures were additionally conducted with paired injections onto the GC-APCI-MS system using split ratio 5 with TMS-derivatized samples and split ratio 100 with TBDMS samples. Retention index standardization was based on *n*-alkanes as described earlier^29^.

### Quantification of aspartic acid

GC-EI-MS chromatograms were recorded at nominal mass resolution, baseline corrected and processed as described previously^29^. Chemical reference compounds and their analytes were picked by manual supervision using TagFinder^31^ and the NIST MS Search 2.0 software (http://chemdata.nist.gov/). Observed experimental mass spectra and retention time indices (RI) were matched to the mass spectral and RI reference collection of the Golm Metabolome Database (GMD)^32^. A######-### identifiers directly relate to GMD entries. Quantification of isotopologues and isotopologue distributions (MIDs) were based on peak apex abundances.

Exact mass of GC-APCI-MS files were internally calibrated based on PFTBA^30^. Files were transcribed into mzXML format using Bruker DataAnalysis and AutomationEngine software (version 4.2). Analytes of GC-APCI-MS files were identified manually based on exact monoisotopic masses, comparison to the paired GC-EI-MS analyses and parallel measurements of metabolite reference compounds. The isotopologue abundances of molecular ions and mass fragments and respective ^13^C labeled MIDs were extracted from each GC-APCI-MS files in a defined chromatographic time range adjusted to each analyte and in a mass range of ± 0.005 mass units using the R packages xcms (version 3.22.0)^33^, MSnbase (version 2.26.0)^34^ and msdata (version 0.40.0)^35^ in RStudio (2023.6.1.524, http://www.posit.co/, R version 4.3.1). Quantification of isotopologue abundances was based on peak area under the peak apex ± 10 scans.

For metabolite quantification, the sum of all isotopologue abundances was used. Metabolite concentration was normalized to internal standard ^13^C_6_-Sorbitol, OD_750_ and sample volume. Molar metabolite concentrations were acquired through parallel analysis of calibration series of non-labeled reference compounds. Mass spectra and chromatogram plots were generated by Bruker DataAnalysis.

### Determination of ^13^mC enrichment

MIDs and resulting ^13^C enrichment calculations of mass features were corrected for natural isotope abundances (NIA) according to their specific molecular formula using RStudio and the IsoCorrectoR package (version 1.18.0)^36^. Correction of tracer impurity was done manually by adjusting IsoCorrectoR results to the specified ^13^C purity of each reference substance. E^13^C and molar concentrations of aspartic acid were used to calculate molar ^13^C concentration at each carbon position of the aspartic acid backbone.

### Calculation of positional ^13^C enrichment

Position-specific ^13^C enrichment of aspartic acid for position 1, 2, 3, and 4 was calculated according to formulas 1, 2, 3, and 4, respectively.

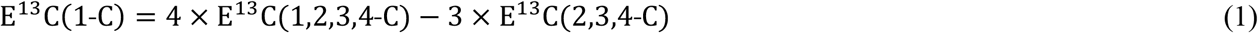

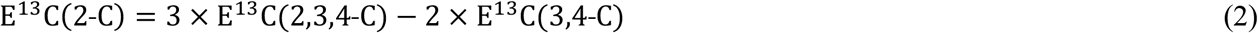

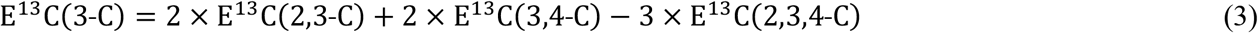

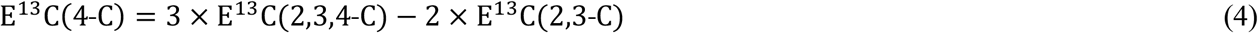

Mass features of aspartic acid analytes from GC-MS analyses that contain required carbon combinations, 1,2,3,4-C, 2,3,4-C, 2,3-C, and 3,4-C and their use for position-specific ^13^C enrichment calculations are reported in the results section.

### Estimation of molar carbon assimilation by sigmoidal curve fitting

Assimilation rates in 1-C and 4-C of aspartate were determined by sigmoidal curve fitting using R Studio and package sicegar (version 0.2.4, threshold_AIC = -10 (default setting))^37^. Absolute positional E^13^C (pmol/OD_750_ × mL^-1^) at 0, 5, 10, 15, 30, 60 and 90 min served as input data. The threshold intensity ratio was set to 0.75 and the maximum allowed intensity at t_0_ to 0. The maximum slope in the sigmoidal plot equals the maximum assimilation rate (pmol/OD_750_ × mL^-1^ × min^-1^) and the time point (min) when maximum assimilation is reached is called “half max”.

## Results and Discussion

### Exploration of GC-MS technology towards ^13^C-positional enrichment analysis of aspartate

To assess *in vivo* activity of PEPC, first we aimed to measure steady state amounts of OAA, and its products malate or aspartate. Malate and aspartate were reliably quantified as reported before^38, 39^, whereas we never obtained measurements of OAA. To check the detection limit of our methods, we subjected different amounts of OAA to GC-EI-MS and GC-APCI-MS analyses. With both systems, it was impossible to detect OAA below 30 ng injected (Figure S1, analyte 1MEOX 2TMS MP (A147010-101), 1 ng for GCAPCI-MS). This amount corresponds to 3.0 nmol/OD_750_*mL^-1^ of *Synechocystis* cultures sampled at OD_750_=1 with a volume of 10 mL (0.1 nmol/OD_750_*mL^-1^ with GC-APCI-MS). Cellular concentrations of OAA were lower likely due to its immediate conversion to product metabolites. Malate concentration in *Synechocystis* extracts were close to the lower detection limit of our GC-MS systems. Flux studies predicted a major flux from OAA to aspartate with minor fluxes to malate and citrate^22^. Therefore, we decided to use positional labelling information from aspartate as proxy of Ci assimilation modes in *Synechocystis*.

To identify the positional E^13^C information in the aspartate carbon backbone, we analyzed in the first step the fragmentation patterns of aspartic acid derivatized with two different derivatization agents, BSTFA and MTBSTFA, commonly used for GC-MS analysis and leading to TMS (SiC_3_H_9_) and TBDMS (SiC_5_H_15_) modifications at carboxy-, hydroxy-, and amino-, and amide-groups, among others. Identification of aspartic acid derivatives in GC-MS chromatograms was achieved by database matching to retention indices and mass spectra (Golm Metabolome Database, http://gmd.mpimp-golm.mpg.de)^32^ combined with measurements of specific ^13^C-labeled aspartic acid standards. Chromatograms and identified peaks after TMS derivatization measured with GC-EI-MS and GC-APCI-MS are reported in Figure S2. All hydroxy- and carboxy-groups were silylated, but silylation of the amino-group was partial and generated two detectable aspartic acid derivatives, 2TMS and 3TMS or 2TBDMS and 3TBDMS, respectively. The analytes with 3 silyl groups were more abundant in standard measurements or biological extracts and less variable. For BSTFA derivatization, aspartic acid 3TMS was preferably used for quantification. For ^13^C enrichment determination all available analytes were tested, because the proportions of fragment ion isotopologues were not directly affected by variation of analyte quantity.

First, all measured mass features of aspartic acid standard substances were screened for the presence of their carbon backbone. Mass spectra from fully labeled [U-^13^C]aspartic acid in comparison to mass spectra of natural aspartic acid were used to ascertain the number of carbon atoms originating from aspartic acid in each fragment by mass shift analysis (**Figure 1**). Positional labeled [1-^13^C], [2-^13^C], [3-^13^C], [4-^13^C]aspartic acids were used to determine the involved carbon atoms (**Figure 1**). This analysis suggested fragment ions involving carbon combinations 1,2,3,4-C, 2,3,4-C, 1,2-C, 2,3-C and 3,4-C of aspartic acid as indicated (**Figure 1**). In most cases, the same fragment ions as in GC-APCI-MS were detected by GC-EIMS (Figure S3). Differences of fragmentation patterns originated from the ionization modes of the two GC-MS technologies. It was previously shown that GC-APCI-MS can be considered as a soft ionization method^40^. However, fragment ions detected by GC-EI-MS were also observed in GC-APCI-MS due to in-source fragmentation mainly through neutral loss reactions^41^. Fragmentation analyses of aspartic acid 2TMS and 3TBDMS are displayed in Figures S4-5.

**Figure 1.**
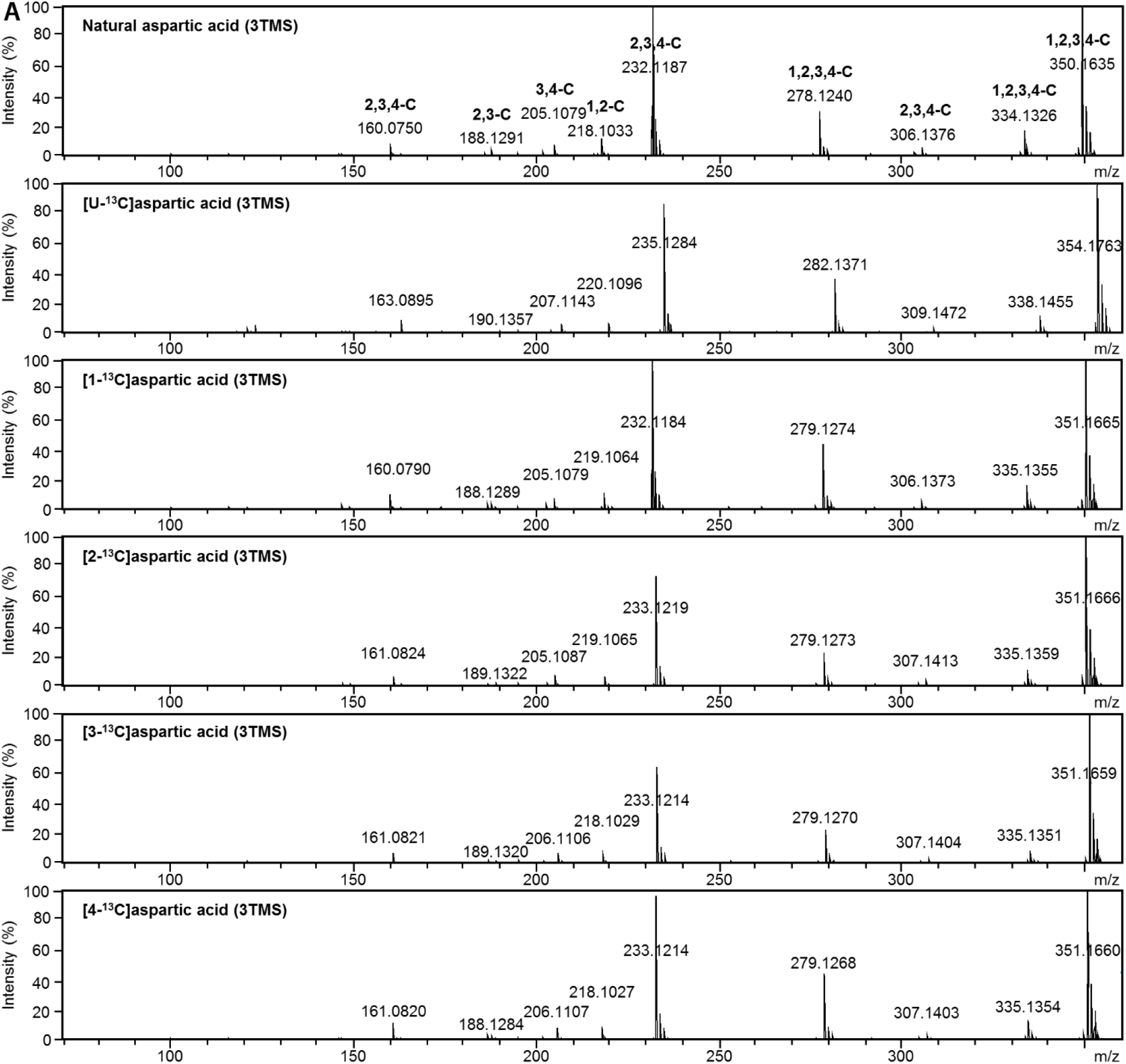
APCI-induced fragmentation of aspartic acid 3TMS. Natural, fully and position-specific ^13^Clabeled aspartic acid was trimethylsilylated (TMS) and separated by GC coupled to atmospheric pressure chemical ionization (APCI) MS. Representative mass spectra of the aspartic acid 3TMS derivatives are displayed. Mass shifts of fragment ions from fully labeled [U-^13^C]aspartic acid indicate the number of included carbon atoms originating from aspartic acid. Mass shifts of fragment ions of positional labeled aspartic acid provide positional information of carbon atoms that are included in a specific fragment. Fragment interpretations are indicated within the mass spectrum of natural aspartic acid. Fragment abundances are normalized to the base peak.

GC-APCI-MS measurements determine the molecular mass of fragments very precisely and thus, were also used to suggest molecular formulas of all observed fragment ions. The molecular formulas are necessary for natural isotope abundance (NIA) correction and determination of E^13^C. Combining information of involved carbon atoms from the aspartic acid backbone and the match of exact mass with predictions from *in silico* fragmentation analyses (Figure S6 for aspartic acid 3TMS), we suggested molecular formulas of most of the fragments appearing in GC-APCI-MS and GC-EI-MS. All identified and interpreted fragments are listed in Table S2 together with their recorded properties.

### Positional purity of fragment identities

Mass features from GC-MS are only providing information with desired specificity if the fragment ions and their MIDs are monitored without interference either by other fragment ions of the respective aspartic acid derivative or by co-eluting compounds of a complex sample matrix. The later interferences may arise with every change of experimental conditions and always need to be considered carefully. To test for the presence of interferences, reference compounds with known E^13^C and their fragments with known molecular formulas can be measured and the deviation of expected from measured E^13^C determined. Previous studies used either precisely labeled standards^21, 25^ or biological samples with controlled stable isotope label^7, 42^. We used the four positionally ^13^C-labeled aspartic acid standards (**Figure 1**) and measured these in paired analyses by GC-EI-MS and GC-APCI-MS. The aim of this validation study was the characterization of the positional purity of fragment ions and the definition of the lower detection limit for E^13^C determinations.

E^13^C accuracy is dependent on the amount of injected substance and can be compromised by chemical or electronic noise at the lower limit or by detector saturation at the upper limit of detection. To find these limits of E^13^C determination, all positional aspartic acid standards were combined equally and then isotopically diluted by natural aspartic acid in different proportions (Table S1). Injections of 1.25 ng up to 250 ng of each mixture were analyzed to cover the abundance-range of our analyses in agreement with the expected biological variation^28, 39^. Most fragment ions showed small absolute value deviations from expected E^13^C when 12.5 ng or more were injected (**Figure 2** A). Fragment ion m/z 232, which was close to base peak abundance, showed an increase of absolute E^13^C deviation for injections > 125 ng and fragment ion abundance > 10^7^ (arbitrary units). This saturation phenomenon was fragment specific and applied to fragment m/z 350, too. Split-injection measurements by GC-MS with adjusted split ratios can be performed when abundance saturation of aspartic acid in a sample should be reached, as was done for the subsequent analyses for fragments m/z 232 and m/z 350 (**Figure 2** B).

**Figure 2.**
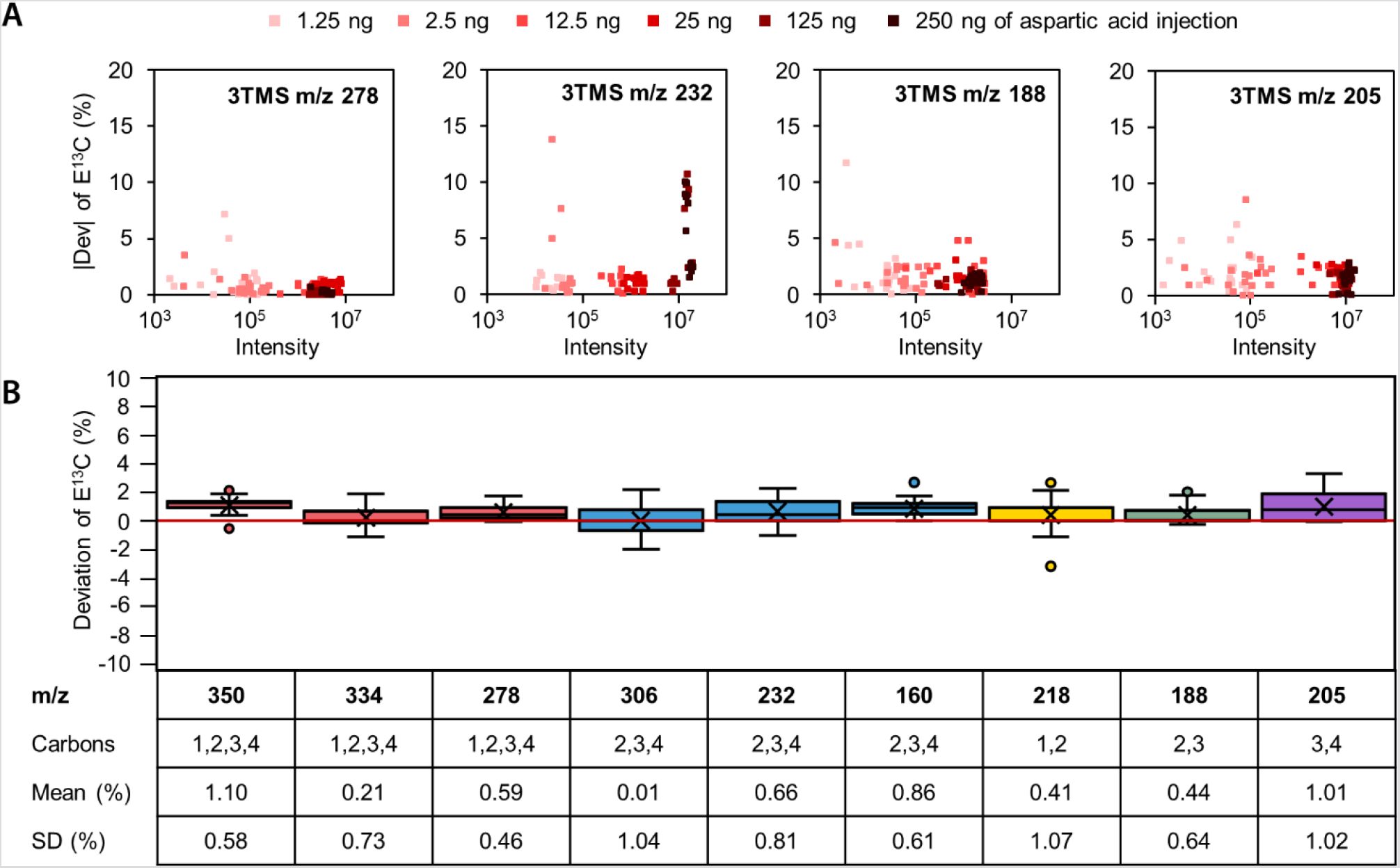
Accuracy and precision of E^13^C determination of aspartic acid 3TMS fragments and adducts by GC-APCI-MS. Equal mixtures of four positional ^13^C-labeled aspartic acid standards in different isotopic dilutions by natural aspartic acid and at different concentrations (Table S1) were measured by GC-APCIMS. Deviations of E^13^C, i.e. measured E^13^C subtracted from expected E^13^C, of the specified fragment ions were analyzed. (A) E^13^C absolute value deviations of selected fragments depended on fragment specific abundances, i.e. the sum of all isotopologue abundances, and amounts of injected aspartic acid. Most fragments provided accurate enrichment information with injections > 12.5 ng aspartic acid. (B) Box-plot representation with means, and standard deviations (SD) of E^13^C deviations from selected fragments defined by nominal mass to charge ratio (m/z). 36 different mixtures were analyzed by 4 technical replicates injecting 25 ng aspartic acid. Indicated mean deviations and SD were determined as parameters of accuracy and precision, respectively. All fragments were analyzed by GC-MS measurements in split-less mode, except fragments m/z 350 and m/z 232 that were acquired by split mode injections at split ratio 1:5 to avoid interference by detector saturation.

To determine positional accuracy of E^13^C information of fragments, the positional ^13^C-labeled aspartic acid standards were mixed with natural aspartic acid in different ratios (Table S1). Sample injection corresponded to optimized 25 ng of aspartic acid (i.e., the sum of labeled and non-labeled aspartic acid). E^13^C was determined using the predicted molecular formulas (Table S2). The deviation of the determined E^13^C from the expected E^13^C was calculated (**Figure 2** B). We used the mean deviation and the SD of the mean deviation as quality parameters for accuracy and preciseness, respectively, with low values indicating high accuracy and preciseness. For aspartic acid 3TMS analyzed by GC-APCI-MS, all fragment ions had mean deviations as well as SDs of the mean deviation less than 1.1%. Accuracy and precision are in the range of deviations found before when analyzing aspartic acid standards by GC-MS/MS for the determination of positional E^13^C^25^. The comparably high deviation of fragment ion m/z 350, which is the [M+H]^+^ adduct ion, originated from the presence of the molecular ion [M]^+^ with overlapping MIDs. If no other fragment ions comprising the whole carbon backbone should be available, the E^13^C of m/z 350 could be corrected for the overlaid MID of m/z 349 using the previously reported correction function CorMID^43^.

In the analyzed injection range (**Figure 2** A), detector saturation was only observed for GC-APCI-MS, not for GC-EI-MS (Figure S7). Overall, the deviation of E^13^C was higher when using GC-EI-MS, likely due to the lower mass resolution of the instrument and thus, lacking separation of fragment ions and isotopologues from overlapping interferences. Acceptable fragment ions from these paired analyses only covered carbon combinations 1,2,3,4-C, 2,3,4-C and 1,2-C. These measurements from aspartic acid 3TMS are insufficient to calculate E^13^C of all single C-positions of aspartic acid. Additionally, GC-EI-MS required higher concentrated samples as the low abundant fragment ions had improved accuracy when injections were higher than 125 ng. Aspartic acid 2TMS as an alternative but lower abundant option is less fragmented compared to aspartic acid 3TMS. When monitored by GC-APCI-MS, potential useful fragment ions of aspartic acid 2TMS again covered only carbon combinations 1,2,3,4-C, 2,3,4-C and 1,2-C (Figure S8). However, aspartic acid 3TMS measured by GC-APCI-MS provided fragments that represented 1,2,3,4-C, 2,3,4-C, 2,3-C, and 3,4-C and thereby the potential to monitor all C positions of aspartic acid. Moreover, E^13^C information of aspartic acid 3TMS and 2TMS can be combined within same samples when both derivatives are detectable.

Aspartic acid 3TBDMS fragmentation by GC-APCI-MS or by GC-EI-MS alone did not cover all fragment ion combinations necessary to calculate all positional E^13^C (Figure S9). Aspartic acid 2TBDMS was only detectable by GC-APCI-MS.

All fragment ions detected by this study through GC-EI-MS and/ or GC-APCI-MS analyses are listed and characterized by Table S2. With standard mixture measurements, we confirmed suggested molecular formulas and excluded presence of interfering fragment ions from the aspartic acid analytes, as well as from laboratory contaminations that are caused by impurities of the materials, solvents and reagents required for GC-MS analysis. However, the background and sample matrix of complex biological extracts will differ. Both need to be considered and interferences excluded by non-sample controls and non-labeled samples generated under the conditions of a stable isotope pulse labelling experiment. The minimum validation requirement should be that NIA correction of the used mass features from extracts of non-labeled samples approximate E^13^C = 0. Fragment ions should be chosen based on purity, i.e. the absence of overlaying interferences, and an abundance within the optimized limits of E^13^C quantification. These ranges can be characterized using non-labeled biological samples that are tested at increasing injection amounts as exemplified (**Figure 2** A). Saturation at the upper limit can be compensated by reanalysis in split injection mode or if necessary by dilution of the chemically derivatized sample by silylation reagent.

### Calculation of positional E^13^C

Positional E^13^C can be calculated using linear equations combining information of different fragment ions. We developed equations that subtract E^13^C among fragment ions, e.g. subtracting E^13^C of fragment A with a smaller number of C-atoms from E^13^C of fragment B with a larger number of C-atoms, weighted by the number of C-atoms present in these fragments. The resulting E^13^C relates to the C-atom(s) that are in this example present in fragment ion B but not in A. The equations (1), (2), (3), and (4) allow to calculate all positional E^13^C based on available fragment ions of aspartic acid 3TMS that represent 1,2,3,4-C, 2,3,4-C, 2,3-C, and 3,4-C. No two fragment ions were available that allowed to calculate E^13^C of 3-C. In this case, we used E^13^C information of three fragment ions. We developed this strategy because fragment ions that represent single C-atoms are rare and typically low abundant within *in source* fragmentation spectra. Direct means of positional measurements were not available in the case of aspartic acid TMS analytes (**Figure 2**; Table S2; Figure S7-S8).

Accuracy and precision of C-positional E^13^C calculations was tested using the standard measurements introduced in the previous section. Accuracy of positional E^13^C was high for all positions and in the range of the direct E^13^C measurements of fragment ions (**Figure 2** B) when analyzing aspartic acid 3TMS by GC-APCI-MS (**Figure 3**). As was expected SD of E^13^C deviation increased due to error propagation. This technical error of positional E^13^C determination was far below biological variation demonstrated later in our studies (see below) and was deemed acceptable for interpretations of biological data. All alternatives to determine positional E^13^C by GC-APCI-MS and TMS derivatization are summarized by Figure S10. Aspartic acid 2TMS was considered for this purpose but contributed only alternative fragment ions representing 1,2,3,4-C and 2,3,4-C. Alternative calculations of positional E^13^C can be used as internal sanity checks provided these measurements are not subject to specific interferences.

**Figure 3.**
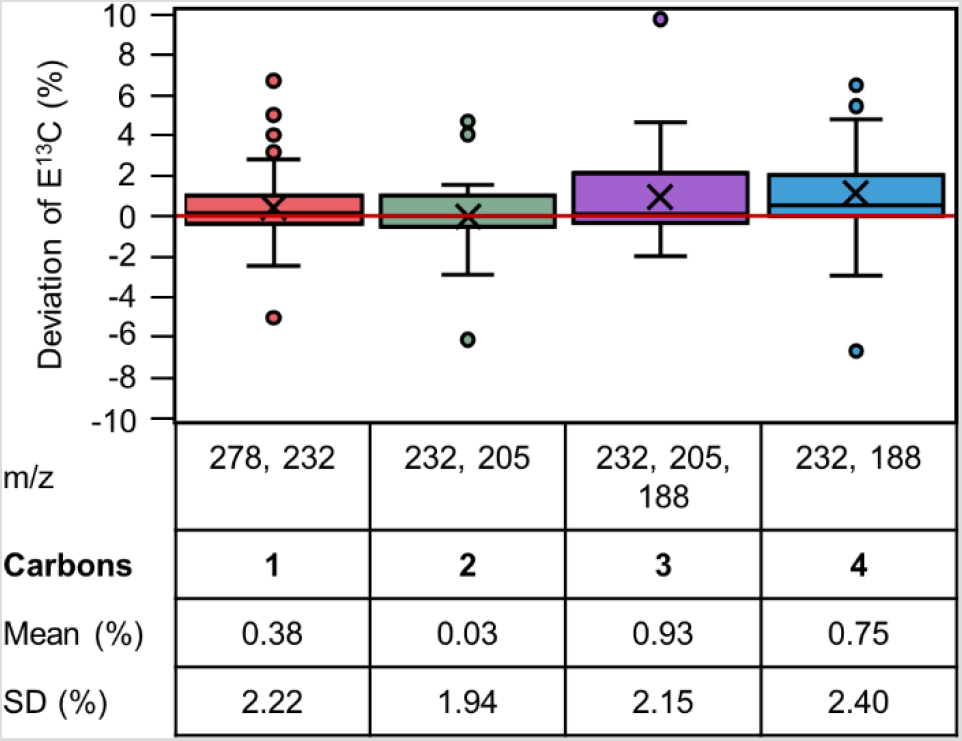
Positional E^13^C accuracy and precision from aspartic acid 3TMS determined by GC-APCI-MS. Positional E^13^C was calculated applying equations (1), (2), (3) and (4) to E^13^C of the indicated fragment ions (m/z). Distribution analyses of E^13^C deviations, i.e. calculated E^13^C subtracted from expected, is displayed across 36 different mixtures with 4 technical replicates (cf. Legend to Figure 2). Amount of total aspartic acid injected equaled 25 ng. E^13^C can be determined at all positions with accuracy (mean) < 1% and precision (SD) < 2.5%. E^13^C of fragment ion m/z 232 was determined by split measurements at split ratio 1:5. E^13^Cs of all other fragment ions were determined by analyses in split-less mode.

GC-EI-MS did allow to determine E^13^C of 3 C-positions, but not all with equal accuracy (Figure S11). E^13^C deviations of 1-C and 2-C calculations were deemed acceptable using optimal alternatives of calculation, whereas 4-C E^13^C was consistently underestimated (Figure S11 B) due to MID interference of fragment ion m/z 188.

Using TBDMS derivatization combined with GC-APCI-MS measurements, we determined positional E^13^C of all positions of aspartic acid with acceptable accuracy but only by combining information of the two analytes, aspartic acid 3TBDMS and aspartic acid 2TBDMS (Figure S12). Precision was, however, inferior compared to TMS derivatized analytes. Even the two TBDMS derivatized fragments that directly represented 2-C of aspartic acid had approximately equal SD compared to the calculations based on TMS derivatization (**Figure 3**; Figure S12 B).

In general, accuracy and precision of positional E^13^C determinations by *in source* mass fragmentations depended on the choice and number of required fragment ions. Considering potential error propagation by the combinatorial calculations, we argued that fragment ions with least mean E^13^C deviation and smallest SD will provide the best positional E^13^C information (**Figure 3**). For biological applications, we suggest to first assess all available E^13^C information from a set of analyzed samples for interferences and inherent technical variability and only then choose the best available combination. As judged by the *Synechocystis* samples of this study, we found the aspartic acid 3TMS analyte superior to the less abundant 2TMS fragment ions with frequent measurements at the lower detection limits and to the TBDMS derivatives with equal or higher SD.

### Determination of PEPC activity *in vivo*

The established C-positional method was combined with dynamic ^13^CO_2_ labelling of *Synechocystis* to estimate PEPC activity and to distinguish this activity from RUBISCO and CBB cycle activities. *Synechocystis* WT was cultivated with high CO_2_ supply (5%) and either in constant light or day/night cycles where ^13^CO_2_ labelling was started in the dark at the beginning of the night. Positional E^13^Cs were determined as described before by analyzing aspartic acid 3TMS within complex extracts of primary metabolites. In constant light, all positions of aspartate were ^13^C labeled with first detectable E^13^C at 10 min after the ^13^CO_2_ pulse (**Figure 4** A). Our data demonstrate that both, PEPC and RUBISCO, assimilate CO_2_ during the day. E^13^C at 1-C of aspartate originates from direct CO_2_ assimilation by RUBISCO into 1-C of 3PGA. Label at 2-C and 3-C of aspartate and 3PGA result from regeneration of RuBP from 3PGA by the CBB cycle. At beginning of the night, first ^13^C label incorporation into aspartate was detected after 15 min. ^13^C label was exclusively delimited to 4-C of aspartate. We concluded that only PEPC was active in the dark and RUBISCO inactive. Our experiments proved previous expectations of PEPC and RUBISCO activity during day and night phases and confirmed that our method indeed distinguished both enzyme activities that contribute to aspartate biosynthesis. We did not detect transfer of ^13^C between carbon positions 1-C and 4-C of aspartate during the night. Such a transfer is potentially conceivable via OAA conversion into malate and a malate to fumarate equilibrium catalyzed by fumarase^20, 44^. The symmetric fumarate molecule allows position-swapping between 1-C and 4-C and may redistribute ^13^C label, if the reversed reaction sequence from fumarate to OAA should be active. *In vitro* studies, however, showed that the K_m_ for the generation of malate from fumarate is higher than K_m_ of the reverse reaction^44^, and a 5-fold higher ^13^C atom number, i.e. μmol (^13^C) * g-dry cell weight^-1^, was present in malate than in fumarate *in vivo*^18^. These studies are indicating that the fumarase acts preferentially unidirectional. Furthermore, flux balance analyses showed that the TCA cycle is operated in the reductive direction with no cyclic flux during the day. This operation-mode of the TCA cycle prevents shuffling of ^13^C label between C-positions^22, 45^. Even though a low cyclic flux was predicted during the night^45^, no redistribution of label between 4-C and 1-C was detected in our studies. Therefore, we concluded that E^13^C at 4-C of aspartate is a specific proxy of PEPC mediated carbon assimilation.

**Figure 4.**
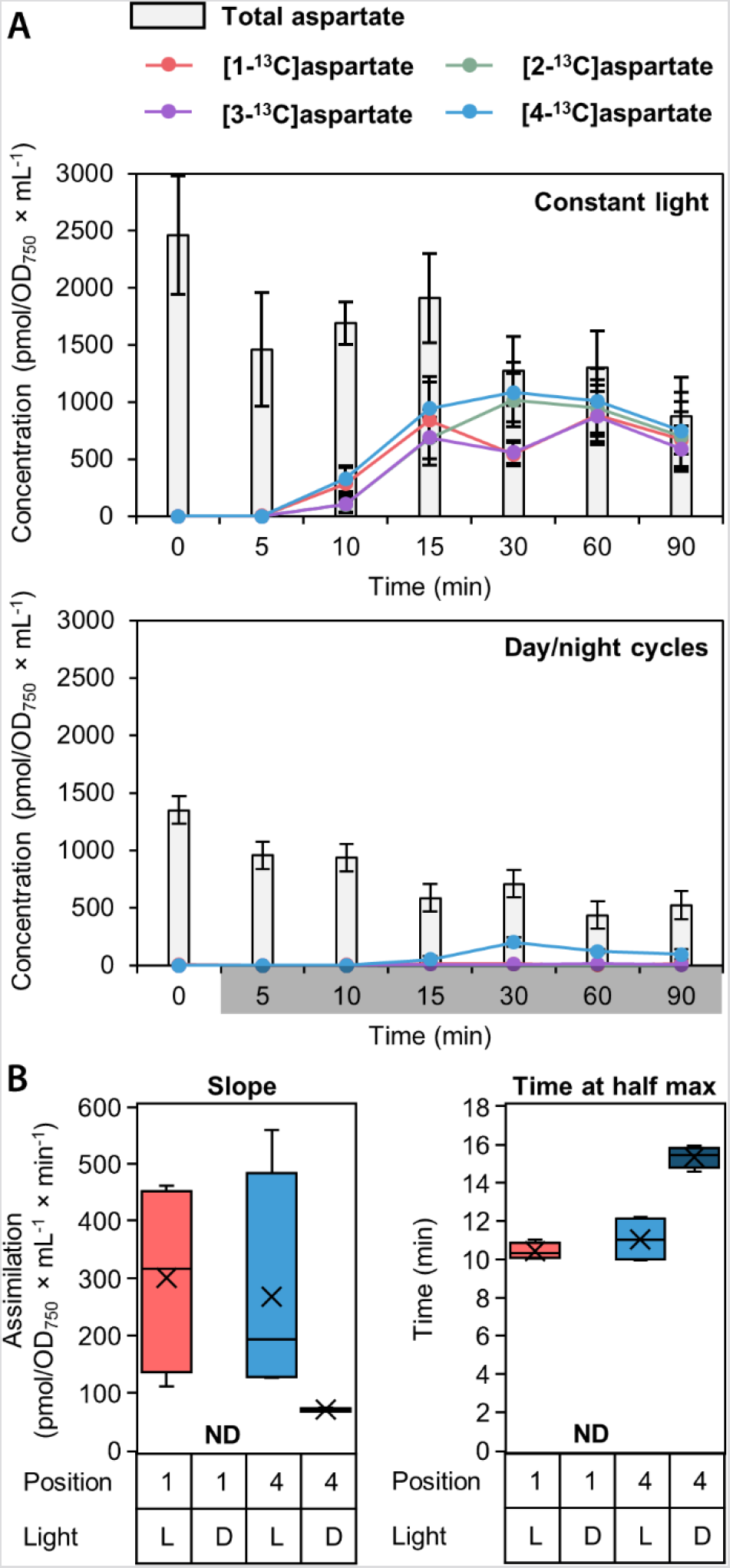
Determination of position-specific ^13^C labelling of aspartate in *Synechocystis* sp. PCC 6803 during the day compared to the night. *Synechocystis* cells were cultivated photoautotrophically with 5% CO_2_-enriched air. Cells were either cultivated in constant light (CL) or a 12 h:12 h day/night photoperiod (DN). For DN, the ^13^CO_2_ labelling pulse was applied after transition to night. Samples were taken 5 to 90 min after the labelling pulse. Samples were analyzed by GC-MS. (A) Absolute aspartate concentrations (pmol/OD_750_ × mL^-1^) were determined (grey bars). Position-specific E^13^C was multiplied with absolute aspartate concentrations (points + lines). Data points represent the mean of 4 biological replicates ± SD. During the day, both enzymes are active. All carbon positions of aspartate assimilate ^13^C. In the night, only PEPC is active demonstrated by ^13^C assimilation into position 4-C. (B) To estimate molar carbon-assimilation rates catalyzed by PEPC and RUBISCO, assimilation curves were fitted sigmoidal (Figure S13). Maximum carbon assimilation (slope) and the timepoint of maximum assimilation (time at half max) are displayed for position 1-C and 4-C in the light (L) and in the dark (D).

*In vivo* PEPC activity depends on enzyme amount per cell, the inherent specific enzyme activity, the regulatory state, and availability of substrates (PEP and HCO_3_^-^). To characterize the CO_2_ assimilation further, molar concentrations, i.e. pmol of total, ^13^C labeled, or non-labeled carbon atoms × OD_750_^-1^ × mL^-1^ were calculated by quantifications of aspartate amounts using GC-EI-MS. Quantifications of E^13^C by GCAPCI-MS and of molar amounts by GC-EI-MS were paired by sample and allowed quantifications of ^13^C labeled molar amounts at each C-position of aspartate across our dynamic ^13^CO_2_ labelling experiments (**Figure 4** A). The resulting molar time series were subjected to sigmoidal curve fitting to estimate the maximal C-assimilation rate into carbon positions of aspartate. For this purpose, we determined the maximum slope of the fitted sigmoidal functions over time (Figure S13). The assimilation rates by RUBISCO into 1-C and by PEPC into 4-C of aspartate during the day were similar, indicating that C-assimilation into aspartate was coordinated and balanced between both enzymes (**Figure 4** B). The variation between the biological replicate time series was caused by light-dependent variation of aspartate concentration measurements rather than by SD of E^13^C (**Figure 4** A).

To interpret the positional carbon assimilation rates into aspartate via RUBISCO, i.e. 1-C of 3PGA, and the CBB cycle, i.e. 2-C and 3-C of 3PGA through regeneration of RuBP, dilution of 3PGA by anaplerotic reactions from non-labeled carbohydrate sources needs to be considered^46^. Likewise, OAA can be generated from extant non-labeled malate. Therefore, we consider assimilation rates into 1-C and 4-C of aspartate as proxies of enzyme activities that are not directly comparable between positions due to different metabolic distances from the respective enzyme activities. However, comparison of assimilation rates into the same position can be compared between different conditions, strains or mutants. We demonstrated that the assimilation rate of PEPC is lower during the night than during the day, and, as was expected, RUBISCO assimilation was not detectable in darkness. The time point of maximal C positional assimilation is a second characteristic of the monitored enzyme activities. During the night, more time was needed to reach maximal assimilation rates (**Figure 4** B). This observation was also in agreement with previous knowledge. The prolonged time until maximum labelling rates is likely due to reduced CO_2_ uptake into *Synechocystis* cells during the night as Ci uptake systems are known to be inactivated in darkness^47^.

## Conlusions

Positional E^13^C analysis of aspartate by GC-MS provides positional E^13^C and accurate molar quantifications simultaneously. Moreover, GC-MS requires less sample material compared to ^13^C positional NMR analysis. Resolving the contribution of different reactions to the carbon backbone of aspartate helps to clarify metabolic routes and allows specification and higher precision of future flux analyses. In photoautotrophic organisms, such as cyanobacteria, Ci assimilation by RUBISCO, the recycling process of RuBP via the CBB cycle, and Ci assimilation by PEPC can be distinguished by the method established in this study. These technological features will be especially helpful when all C-assimilating reactions are active, e.g. in photoautotrophic growth conditions. Application fields cover the determination of changes in PEPC activity under changing environmental conditions and among different genetic backgrounds or mutants. The analysis pipeline we established in this study can be extended to the various metabolites that are simultaneously detectable by GC-MS and is only constrained by the compound specific *in source* fragmentation reactions.

### Associated Content Supporting Information

The Supporting Information is available free of charge on the ACS Publications website.

Standard mixtures, oxaloacetate detection, mass spectra, fragment characterization of TMS and TBDMS analytes, *in silico* fragmentation, accuracy and precision of TMS and TBDMS analytes, positional E^13^C of TMS and TBDMS analytes, sigmoidal curve fitting (PDF)

## Author Information

## Author Contributions

JK, MH and LW designed the study. LW conducted standard mixture experiments. LW and AE analyzed the data. LW and YR conducted ^13^CO_2_ labelling experiments. LW and JK wrote the manuscript. All authors edited the manuscript. All authors have given approval to the final version of the manuscript.

## Acknowledgements

We acknowledge the funding of the German Research Foundation (DFG) in the framework of the research consortium SCyCode (FOR2816; KO 2329/7-2, HA2002/23-2) and support by the Max Planck Society. We thank Prof. Dr. Zoran Nikoloski and Prof. Dr. Elke Dittmann for helpful discussions.

## Notes

### Competing Interest Statement

The authors have declared no competing interest.

