## Supplementary information for "Positional ^13^C Enrichment Analysis of Aspartate by GC-MS to Determine PEPC Activity *In Vivo*"

Figure S 1. Detection limit of oxaloacetate analyzed by GC-EI-MS. 3
Figure S 2. Gas chromatographic separation of aspartic acid TMS derivatives by GC-EI-MS and GC-APCI-MS. 5
Figure S 3. EI-induced fragmentation of 3TMS-derivatized aspartic acid. 6
Figure S 4. EI- and APCI-induced fragmentation of aspartic acid 2TMS. 7
Figure S 5. EI- and APCI-induced fragmentation of aspartic acid 3TBDMS. 8
Figure S 6. *In silico* fragmentation analysis of aspartic acid 3TMS. 10
Figure S 7. Accuracy and precision of $E^{13}C$ determination of aspartic acid (3TMS) by GC-EI-MS. 11
Figure S 8. Accuracy of $E^{13}C$ determination of aspartic acid (2TMS) by GC-APCI-MS. 12
Figure S 9. Accuracy of $E^{13}C$ determination of aspartic acid (3TBDMS) by GC-APCI-MS. 13
Figure S 10. Possibilities to calculate all C-positional $E^{13}C$ of aspartic acid using derivatives 3TMS and 2TMS by GC-APCI-MS. 16
Figure S 11. Options of positional $E^{13}C$ calculations of aspartic acid using GC-EI-MS. 17
Figure S 12. Options of positional $E^{13}C$ determinations of aspartic acid by GC-APCI-MS after TBDMS derivatization. 19
Figure S 13. Sigmoidal curve fitting of aspartate 1-C and 4-C labeling within *Synechocystis* cultures during the day and the night. 20

Table S 1. Composition of standard mixtures. 2
Table S 2. Fragment ion validation of trimethylsilylated and tert-butyldimethylsilylated derivatives of aspartic acid. 14

**Table S 1. Composition of standard mixtures.** Mixtures to determine the lower threshold of $E^{13}C$ detection are shown in green (ID 32-64). Accuracy and precision of $E^{13}C$ were determined by mixtures indicated by yellow underlay, all with 25 ng; (ID 1-31, ID 42-46). Amount injected (ng) refers to the amount of aspartic acid standard after chemical derivatization and injected for GC-MS measurements.

| | | Proportion (%) | | | | | | Amount injected (ng) | | | | | |
|---|---|---|---|---|---|---|---|---|---|---|---|---|---|
| ID | amount (ng) | natural | $U\text{-}^{13}C$ | $1\text{-}^{13}C_1$ | $2\text{-}^{13}C_1$ | $3\text{-}^{13}C_1$ | $4\text{-}^{13}C_1$ | natural | $U\text{-}^{13}C$ | $1\text{-}^{13}C_1$ | $2\text{-}^{13}C_1$ | $3\text{-}^{13}C_1$ | $4\text{-}^{13}C_1$ |
| **1** | 25 | 100 | 0 | 0 | 0 | 0 | 0 | 25 | 0 | 0 | 0 | 0 | 0 |
| **2** | 25 | 0 | 100 | 0 | 0 | 0 | 0 | 0 | 25 | 0 | 0 | 0 | 0 |
| **3** | 25 | 0 | 0 | 100 | 0 | 0 | 0 | 0 | 0 | 25 | 0 | 0 | 0 |
| **4** | 25 | 0 | 0 | 0 | 100 | 0 | 0 | 0 | 0 | 0 | 25 | 0 | 0 |
| **5** | 25 | 0 | 0 | 0 | 0 | 100 | 0 | 0 | 0 | 0 | 0 | 25 | 0 |
| **6** | 25 | 0 | 0 | 0 | 0 | 0 | 100 | 0 | 0 | 0 | 0 | 0 | 25 |
| **7** | 25 | 95 | 0 | 5 | 0 | 0 | 0 | 23.75 | 0 | 1.25 | 0 | 0 | 0 |
| **8** | 25 | 90 | 0 | 10 | 0 | 0 | 0 | 22.5 | 0 | 2.5 | 0 | 0 | 0 |
| **9** | 25 | 50 | 0 | 50 | 0 | 0 | 0 | 12.5 | 0 | 12.5 | 0 | 0 | 0 |
| **10** | 25 | 10 | 0 | 90 | 0 | 0 | 0 | 2.5 | 0 | 22.5 | 0 | 0 | 0 |
| **11** | 25 | 5 | 0 | 95 | 0 | 0 | 0 | 1.25 | 0 | 23.75 | 0 | 0 | 0 |
| **12** | 25 | 95 | 0 | 0 | 5 | 0 | 0 | 23.75 | 0 | 0 | 1.25 | 0 | 0 |
| **13** | 25 | 90 | 0 | 0 | 10 | 0 | 0 | 22.5 | 0 | 0 | 2.5 | 0 | 0 |
| **14** | 25 | 50 | 0 | 0 | 50 | 0 | 0 | 12.5 | 0 | 0 | 12.5 | 0 | 0 |
| **15** | 25 | 10 | 0 | 0 | 90 | 0 | 0 | 2.5 | 0 | 0 | 22.5 | 0 | 0 |
| **16** | 25 | 5 | 0 | 0 | 95 | 0 | 0 | 1.25 | 0 | 0 | 23.75 | 0 | 0 |
| **17** | 25 | 95 | 0 | 0 | 0 | 5 | 0 | 23.75 | 0 | 0 | 0 | 1.25 | 0 |
| **18** | 25 | 90 | 0 | 0 | 0 | 10 | 0 | 22.5 | 0 | 0 | 0 | 2.5 | 0 |
| **19** | 25 | 50 | 0 | 0 | 0 | 50 | 0 | 12.5 | 0 | 0 | 0 | 12.5 | 0 |
| **20** | 25 | 10 | 0 | 0 | 0 | 90 | 0 | 2.5 | 0 | 0 | 0 | 22.5 | 0 |
| **21** | 25 | 5 | 0 | 0 | 0 | 95 | 0 | 1.25 | 0 | 0 | 0 | 23.75 | 0 |
| **22** | 25 | 95 | 0 | 0 | 0 | 0 | 5 | 23.75 | 0 | 0 | 0 | 0 | 1.25 |
| **23** | 25 | 90 | 0 | 0 | 0 | 0 | 10 | 22.5 | 0 | 0 | 0 | 0 | 2.5 |
| **24** | 25 | 50 | 0 | 0 | 0 | 0 | 50 | 12.5 | 0 | 0 | 0 | 0 | 12.5 |
| **25** | 25 | 10 | 0 | 0 | 0 | 0 | 90 | 2.5 | 0 | 0 | 0 | 0 | 22.5 |
| **26** | 25 | 5 | 0 | 0 | 0 | 0 | 95 | 1.25 | 0 | 0 | 0 | 0 | 23.75 |
| **27** | 25 | 0 | 0 | 95 | 0 | 0 | 5 | 0 | 0 | 23.75 | 0 | 0 | 1.25 |
| **28** | 25 | 0 | 0 | 90 | 0 | 0 | 10 | 0 | 0 | 22.5 | 0 | 0 | 2.5 |
| **29** | 25 | 0 | 0 | 50 | 0 | 0 | 50 | 0 | 0 | 12.5 | 0 | 0 | 12.5 |
| **30** | 25 | 0 | 0 | 10 | 0 | 0 | 90 | 0 | 0 | 2.5 | 0 | 0 | 22.5 |
| **31** | 25 | 0 | 0 | 5 | 0 | 0 | 95 | 0 | 0 | 1.25 | 0 | 0 | 23.75 |
| **32** | 250 | 0 | 0 | 25 | 25 | 25 | 25 | 0 | 0 | 62.5 | 62.5 | 62.5 | 62.5 |
| **33** | 250 | 50 | 0 | 12.5 | 12.5 | 12.5 | 12.5 | 125 | 0 | 31.25 | 31.25 | 31.25 | 31.25 |
| **34** | 250 | 80 | 0 | 5 | 5 | 5 | 5 | 200 | 0 | 12.5 | 12.5 | 12.5 | 12.5 |
| **35** | 250 | 90 | 0 | 2.5 | 2.5 | 2.5 | 2.5 | 225 | 0 | 6.25 | 6.25 | 6.25 | 6.25 |
| **36** | 250 | 96 | 0 | 1 | 1 | 1 | 1 | 240 | 0 | 2.5 | 2.5 | 2.5 | 2.5 |
| **37** | 125 | 0 | 0 | 25 | 25 | 25 | 25 | 0 | 0 | 31.25 | 31.25 | 31.25 | 31.25 |
| **38** | 125 | 50 | 0 | 12.5 | 12.5 | 12.5 | 12.5 | 62.5 | 0 | 15.625 | 15.625 | 15.625 | 15.625 |
| **39** | 125 | 80 | 0 | 5 | 5 | 5 | 5 | 100 | 0 | 6.25 | 6.25 | 6.25 | 6.25 |
| **40** | 125 | 90 | 0 | 2.5 | 2.5 | 2.5 | 2.5 | 112.5 | 0 | 3.125 | 3.125 | 3.125 | 3.125 |
| **41** | 125 | 96 | 0 | 1 | 1 | 1 | 1 | 120 | 0 | 1.25 | 1.25 | 1.25 | 1.25 |
| **42** | 25 | 0 | 0 | 25 | 25 | 25 | 25 | 0 | 0 | 6.25 | 6.25 | 6.25 | 6.25 |
| **43** | 25 | 50 | 0 | 12.5 | 12.5 | 12.5 | 12.5 | 12.5 | 0 | 3.125 | 3.125 | 3.125 | 3.125 |
| **44** | 25 | 80 | 0 | 5 | 5 | 5 | 5 | 20 | 0 | 1.25 | 1.25 | 1.25 | 1.25 |
| **45** | 25 | 90 | 0 | 2.5 | 2.5 | 2.5 | 2.5 | 22.5 | 0 | 0.625 | 0.625 | 0.625 | 0.625 |
| **46** | 25 | 96 | 0 | 1 | 1 | 1 | 1 | 24 | 0 | 0.25 | 0.25 | 0.25 | 0.25 |
| **47** | 12.5 | 0 | 0 | 25 | 25 | 25 | 25 | 0 | 0 | 3.125 | 3.125 | 3.125 | 3.125 |
| **48** | 12.5 | 50 | 0 | 12.5 | 12.5 | 12.5 | 12.5 | 6.25 | 0 | 1.5625 | 1.5625 | 1.5625 | 1.5625 |
| **49** | 12.5 | 80 | 0 | 5 | 5 | 5 | 5 | 10 | 0 | 0.625 | 0.625 | 0.625 | 0.625 |
| **50** | 12.5 | 90 | 0 | 2.5 | 2.5 | 2.5 | 2.5 | 11.25 | 0 | 0.3125 | 0.3125 | 0.3125 | 0.3125 |
| **51** | 12.5 | 96 | 0 | 1 | 1 | 1 | 1 | 12 | 0 | 0.125 | 0.125 | 0.125 | 0.125 |
| **52** | 2.5 | 0 | 0 | 25 | 25 | 25 | 25 | 0 | 0 | 0.625 | 0.625 | 0.625 | 0.625 |
| **53** | 2.5 | 50 | 0 | 12.5 | 12.5 | 12.5 | 12.5 | 1.25 | 0 | 0.3125 | 0.3125 | 0.3125 | 0.3125 |
| **54** | 2.5 | 80 | 0 | 5 | 5 | 5 | 5 | 2 | 0 | 0.125 | 0.125 | 0.125 | 0.125 |
| **55** | 2.5 | 90 | 0 | 2.5 | 2.5 | 2.5 | 2.5 | 2.25 | 0 | 0.0625 | 0.0625 | 0.0625 | 0.0625 |
| **56** | 2.5 | 96 | 0 | 1 | 1 | 1 | 1 | 2.4 | 0 | 0.025 | 0.025 | 0.025 | 0.025 |
| **57** | 1.25 | 0 | 0 | 25 | 25 | 25 | 25 | 0 | 0 | 0.3125 | 0.3125 | 0.3125 | 0.3125 |
| **58** | 1.25 | 50 | 0 | 12.5 | 12.5 | 12.5 | 12.5 | 0.625 | 0 | 0.1563 | 0.1563 | 0.1563 | 0.1563 |
| **59** | 1.25 | 80 | 0 | 5 | 5 | 5 | 5 | 1 | 0 | 0.0625 | 0.0625 | 0.0625 | 0.0625 |
| **60** | 1.25 | 90 | 0 | 2.5 | 2.5 | 2.5 | 2.5 | 1.125 | 0 | 0.0313 | 0.0313 | 0.0313 | 0.0313 |
| **61** | 1.25 | 96 | 0 | 1 | 1 | 1 | 1 | 1.2 | 0 | 0.0125 | 0.0125 | 0.0125 | 0.0125 |

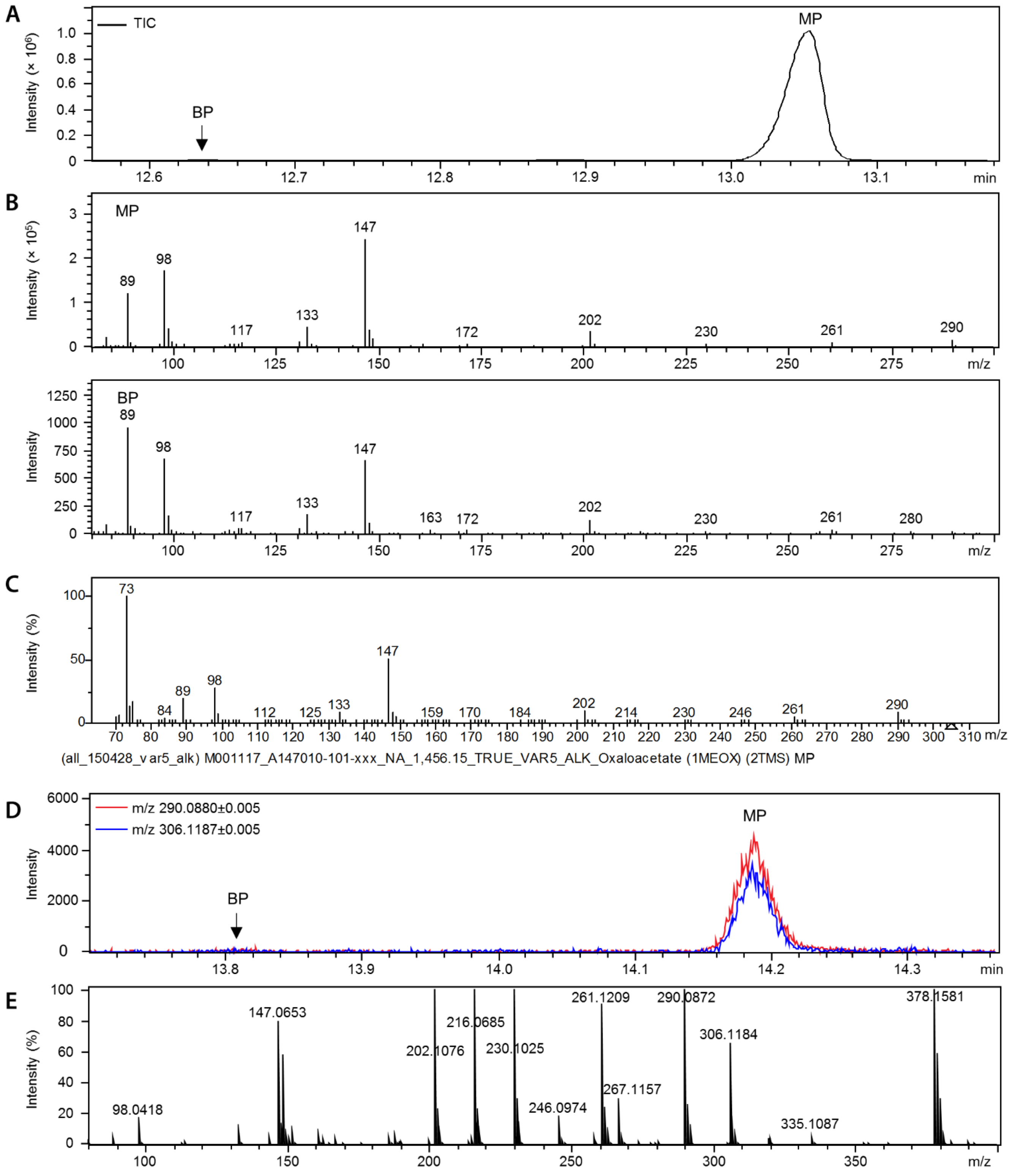


**Figure S 1. Detection limit of oxaloacetate analyzed by GC-EI-MS.** (A, B) Oxaloacetate was methoxyaminated and trimethylsilylated. 30 ng were injected and analyzed by GC-EI-MS. The analytes Oxaloacetate 1MEOX 2TMS MP (A147010-101, main product) and Oxaloacetate 1MEOX 2TMS BP (A142008-101, by-product, indicated by vertical arrow) were detected at chromatographic retention times shown by a total ion chromatogram (A). Determined retention indices (RIs) based on n-alkanes were with 1,454.46 and 1,426.15 close to the reference RIs, 1,456.15 and 1,427.93, respectively, of the Golm Metabolome Database (GMD); http://gmd.mpimp-golm.mpg.de/. No peaks were detected when less than 30 ng were injected. (B) Characteristic mass spectra of Oxaloacetate 1MEOX 2TMS MP (top) and BP (bottom). (C) Mass spectrum of Oxaloacetate 1MEOX 2TMS MP from the GMD database. (D) Oxaloacetate was methoxiaminated and trimethylsilylated. 1 ng was injected and analyzed by GC-APCI-MS. Characteristic mass features, namely m/z 306.1187, i.e. the $H^+$ adduct and m/z 290.0880, i.e. $CH_4$ elimination product from the $H^+$ adduct, are shown. Peaks corresponding to Oxaloacetate 1MEOX 2TMS MP and BP (indicated by vertical arrow) were detected. No peaks were observed by GC-APCI-MS when less than 1 ng was injected. (E) Mass spectrum of Oxaloacetate 1MEOX 2TMS MP measured by GC-APCI-MS with an injected amount of 5 ng.

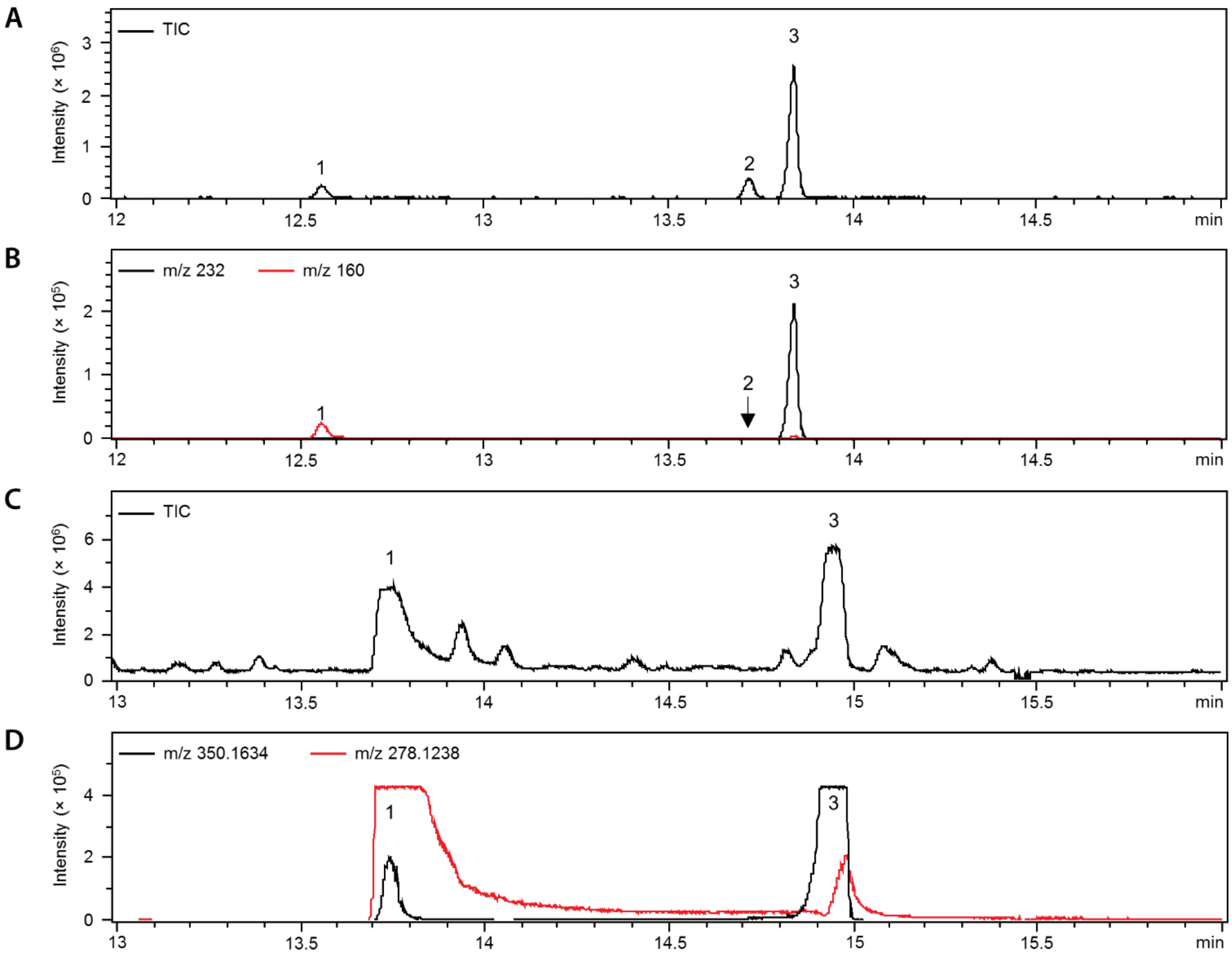


**Figure S 2. Gas chromatographic separation of aspartic acid TMS derivatives by GC-EI-MS and GC-APCI-MS.** Natural aspartic acid was trimethylsilylated (TMS) and 25 ng of aspartic acid were subjected to GC-EI-MS (A, B) and GC-APCI-MS (C, D). Two aspartic acid analytes are detectable by both systems: Aspartic acid 2TMS (peak 1) and aspartic acid 3TMS (peak 3). Peak 2 indicated within the EI total ion chromatograms (TIC) represents *n*-pentadecane used next to other *n*-alkanes for retention index standardization (position indicated by arrow in extracted ion chromatograms in B). Intensity of characteristic masses m/z 160 and m/z 278.1238 for aspartic acid 2TMS and m/z 232 and m/z 350.1634 for aspartic acid 3TMS are displayed in B and D. Please note that aspartic acid 2TMS and 3TMS analytes in APCI are overloaded (C, D) at 25 ng injected. Intermolecular reactions may occur *in source*. Differences in elution time between GC-EI-MS and GC-APCI-MS are due to different column ages and chromatographic differences at the end of the capillary columns with transfer into the alternative ionization units; in case of GC-EI-MS the capillary column ends into a high vacuum for electron impact ionization; in case of GC-APCI-MS the GC effluent is transferred into an atmospheric pressure environment.

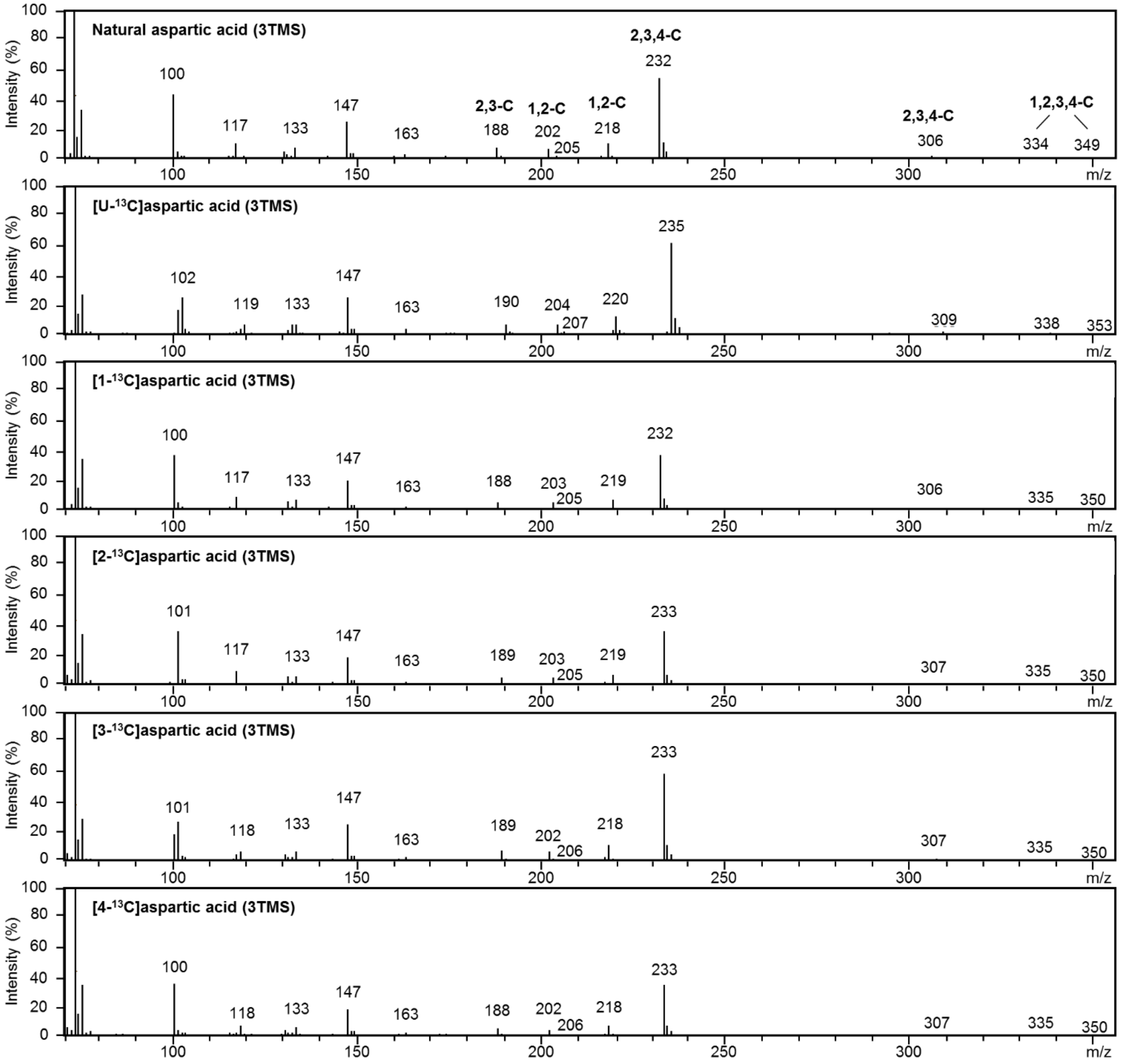


**Figure S 3. EI-induced fragmentation of 3TMS-derivatized aspartic acid.** Natural, fully, and position-specific $^{13}C$-labeled aspartic acids were trimethylsilylated (TMS) and separated by GC coupled to electron ionization (EI) MS. The mass spectra of the aspartic acid 3TMS derivative are displayed. Mass shifts of fragment ions within the mass spectrum of fully labeled [U-$^{13}C$]aspartic acid indicate the number of included carbon atoms originating from aspartic acid. Mass shifts of fragment ions of positional labeled aspartic acid provide information on carbon atoms that are included in a specific fragment as indicated within the natural aspartic acid mass spectrum (top). Intensities (%) are normalized to the base peak.

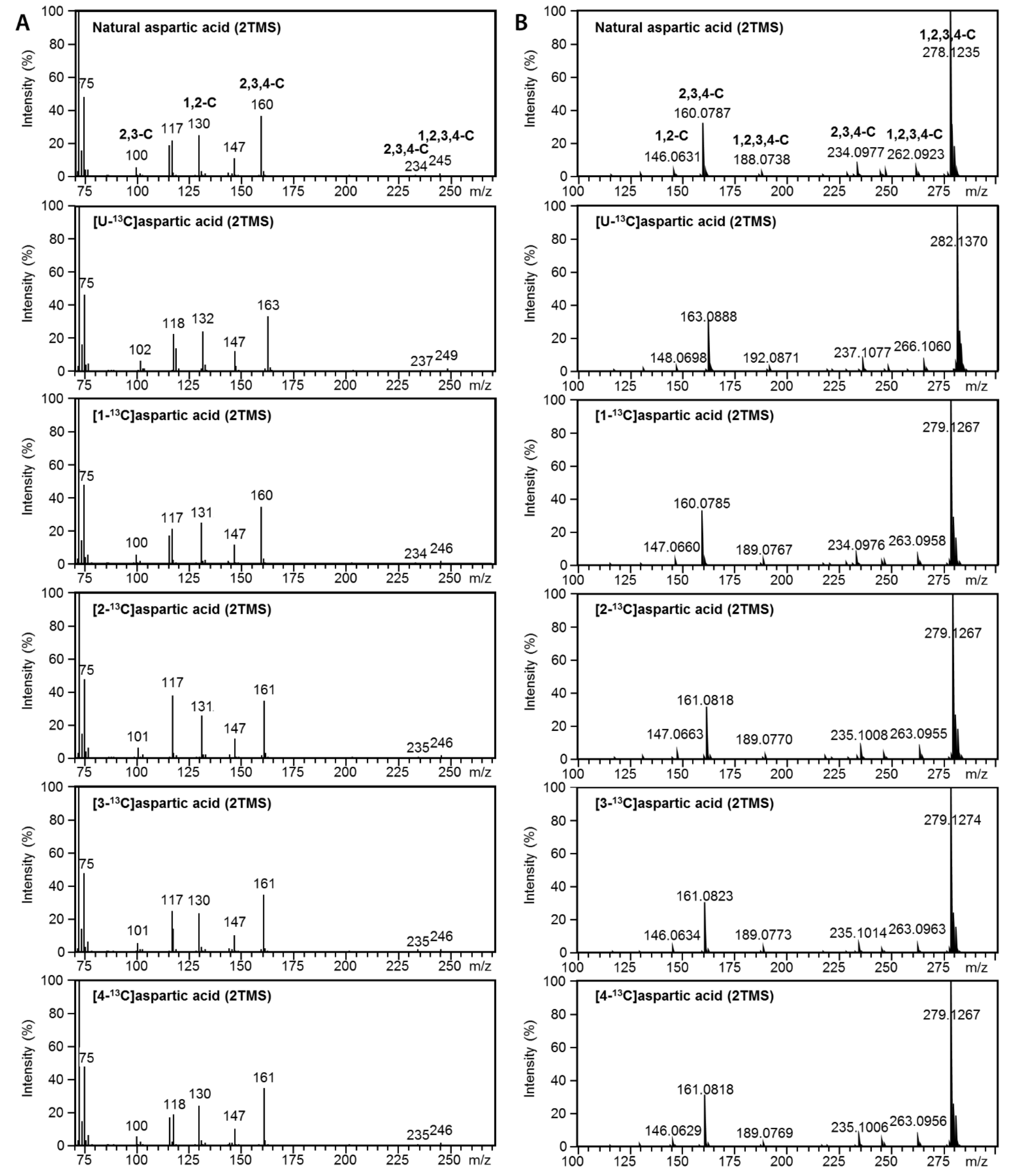


**Figure S 4. EI- and APCI-induced fragmentation of aspartic acid 2TMS.** Natural, fully, and position-specific $^{13}C$-labeled aspartic acids were trimethylsilylated (TMS) and separated by GC coupled to either electron impact ionization (EI) MS (A) or atmospheric pressure chemical ionization (APCI) MS (B). Mass spectra of the aspartic acid 2TMS derivatives are displayed. Mass shifts of fragment ions within the mass spectrum of fully labeled [U-$^{13}C$]aspartic acid indicate the number of included carbon atoms originating from aspartic acid. Mass shifts of fragment ions from positional labeled aspartic acid provide information on the carbon atoms that are included in a specific fragment as indicated within the natural aspartic acid mass spectrum (top). Intensities (%) are normalized to the base peak.

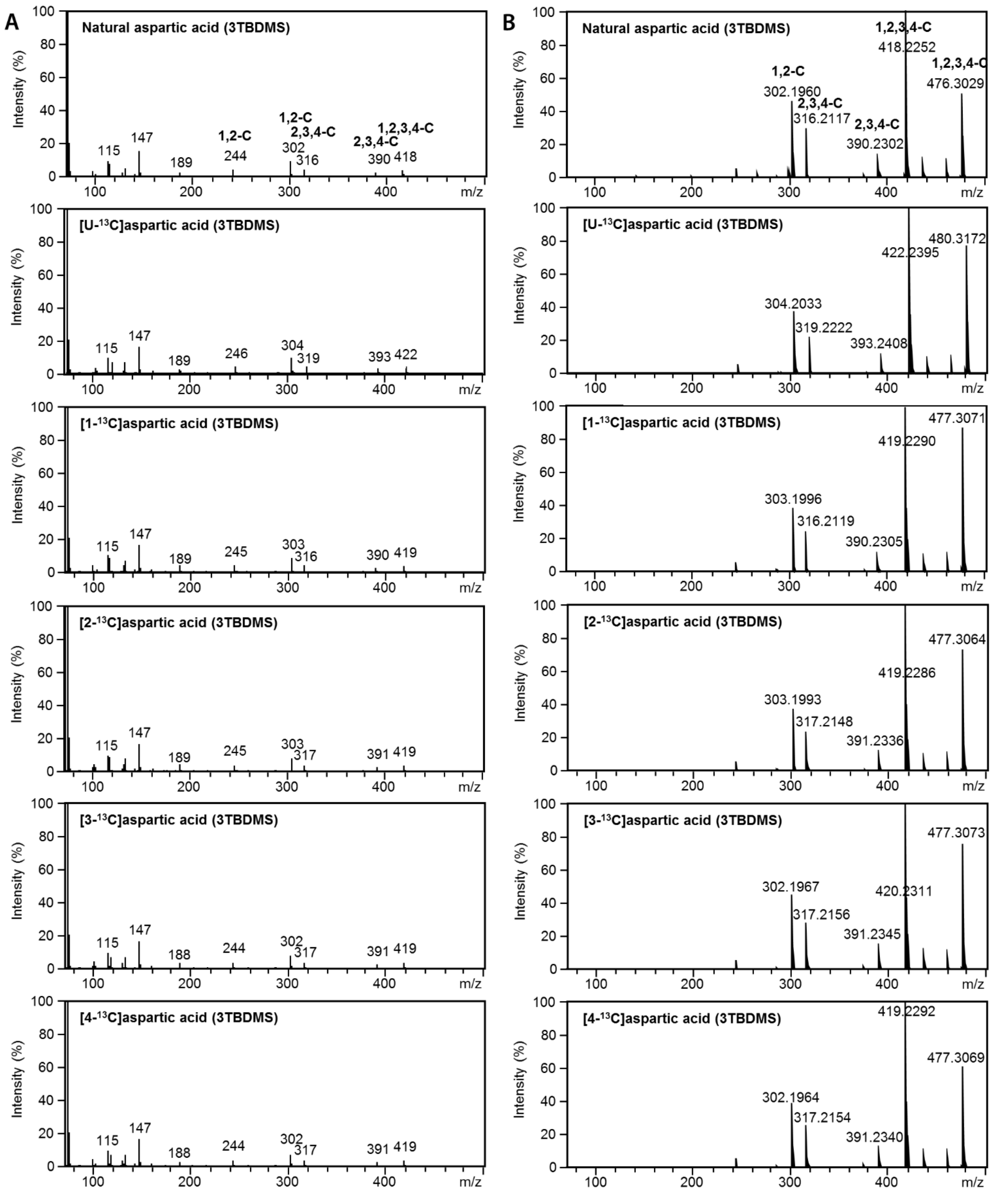


**Figure S 5. EI- and APCI-induced fragmentation of aspartic acid 3TBDMS.** Natural, fully, and position-specific $^{13}C$-labeled aspartic acids were tert-butyldimethylsilylated (TBDMS) and separated by GC coupled to either electron impact ionization (EI) MS (A) or atmospheric pressure chemical ionization (APCI) MS (B). Mass spectra of the aspartic acid 3TBDMS derivatives are displayed. Mass shifts of fragment ions within the mas spectrum of fully labeled[U-$^{13}C$]aspartic acid indicate the number of included carbon atoms originating from aspartic acid. Mass shifts of fragment ions from positional labeled aspartic acid provide information on the carbons atoms that are included in a specific fragment as indicated within the natural aspartic acid mass spectrum (top). Intensities (%) are normalized to the base peak.

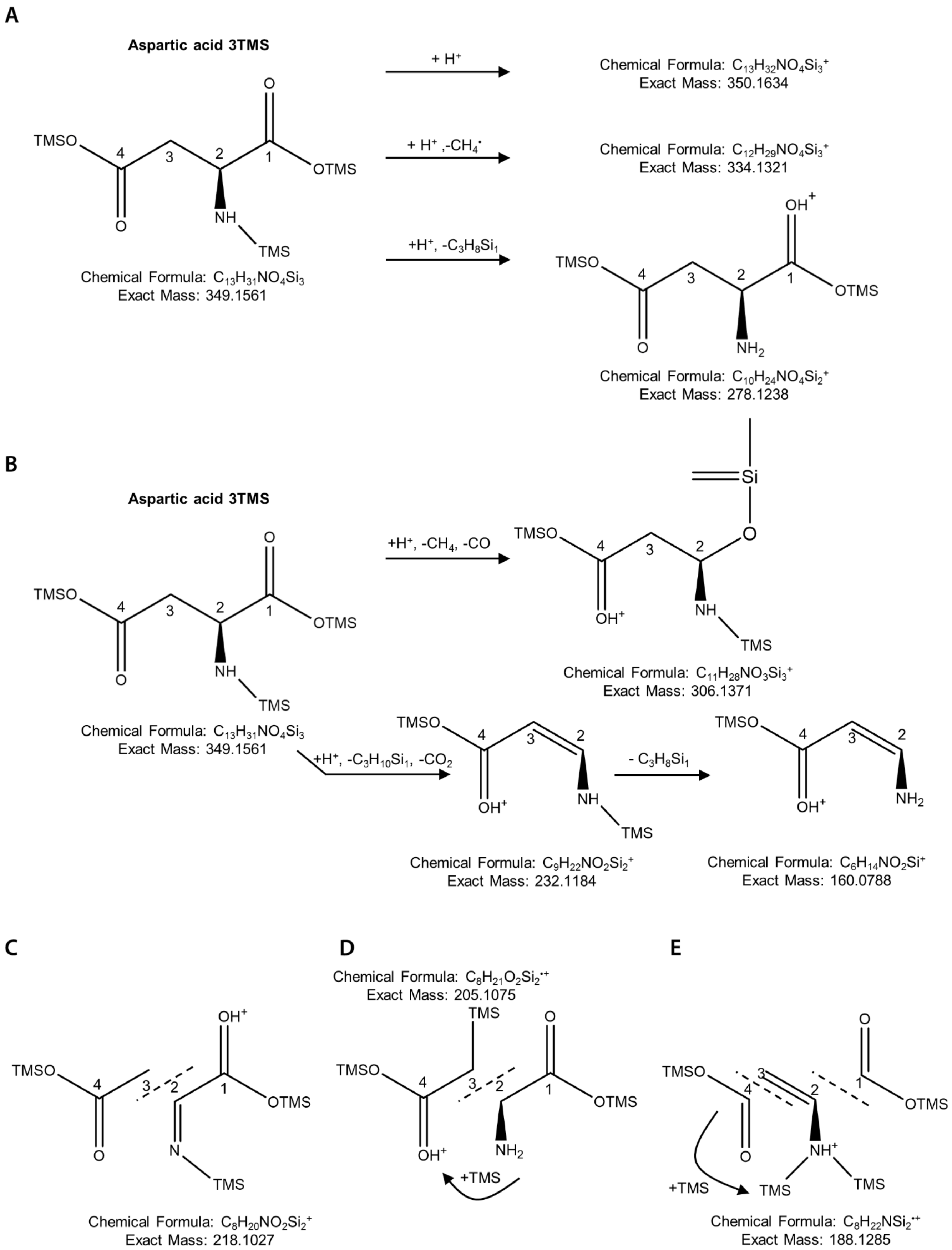
A
Aspartic acid 3TMS
TMSO
OTMS
NH
TMS
Chemical Formula: $C_{13}H_{31}NO_4Si_3$
Exact Mass: 349.1561
+ $H^+$
Chemical Formula: $C_{13}H_{32}NO_4Si_3^+$
Exact Mass: 350.1634
+ $H^+$ ,-$CH_4^{\cdot}$
Chemical Formula: $C_{12}H_{29}NO_4Si_3^+$
Exact Mass: 334.1321
+$H^+$, -$C_3H_8Si_1$
Chemical Formula: $C_{10}H_{24}NO_4Si_2^+$
Exact Mass: 278.1238
B
Aspartic acid 3TMS
Chemical Formula: $C_{13}H_{31}NO_4Si_3$
Exact Mass: 349.1561
+$H^+$, -$CH_4$, -CO
Chemical Formula: $C_{11}H_{28}NO_3Si_3^+$
Exact Mass: 306.1371
+$H^+$, -$C_3H_{10}Si_1$, -$CO_2$
Chemical Formula: $C_9H_{22}NO_2Si_2^+$
Exact Mass: 232.1184
- $C_3H_8Si_1$
Chemical Formula: $C_6H_{14}NO_2Si^+$
Exact Mass: 160.0788
C
Chemical Formula: $C_8H_{20}NO_2Si_2^+$
Exact Mass: 218.1027
D
Chemical Formula: $C_8H_{21}O_2Si_2^{\cdot+}$
Exact Mass: 205.1075
+TMS
E
+TMS
Chemical Formula: $C_8H_{22}NSi_2^{\cdot+}$
Exact Mass: 188.1285

**Figure S 6. *In silico* fragmentation analysis of aspartic acid 3TMS.** Aspartic acid 3TMS fragmentation during GC-APCI-MS was predicted by *in silico* analyses assuming initial protonation, sequential neutral eliminations, and rearrangement or transfer reactions. (A) Different fragment ions with the complete carbon backbone of aspartic acid (1,2,3,4-C) are formed through eliminations of methyl- and/or TMS-groups after proton adduct formation. These moieties originate from the chemical derivatization reagent. (B) Fragment ions including 2,3,4-C of the aspartic acid carbon-backbone are interpreted as result of cleavage between 1-C and 2-C with progressive eliminations of the TMS-moieties. (C) Fragment ion m/z 218 (1,2-C) is thought to arise from cleavage of the aspartic acid molecule between 2-C and 3-C. (D) Likely, rearrangement or transfer of one TMS group to the 3,4-C fragment leads to the formation of fragment ion m/z 205. (E) Cleavages between 1-C/ 2-C and 3-C/ 4-C with rearrangement or transfer of one TMS group to the amino group can explain emergence of fragment ion m/z 188 (2,3-C). Suggested structures are exemplary and may represent one of multiple possible isomers.

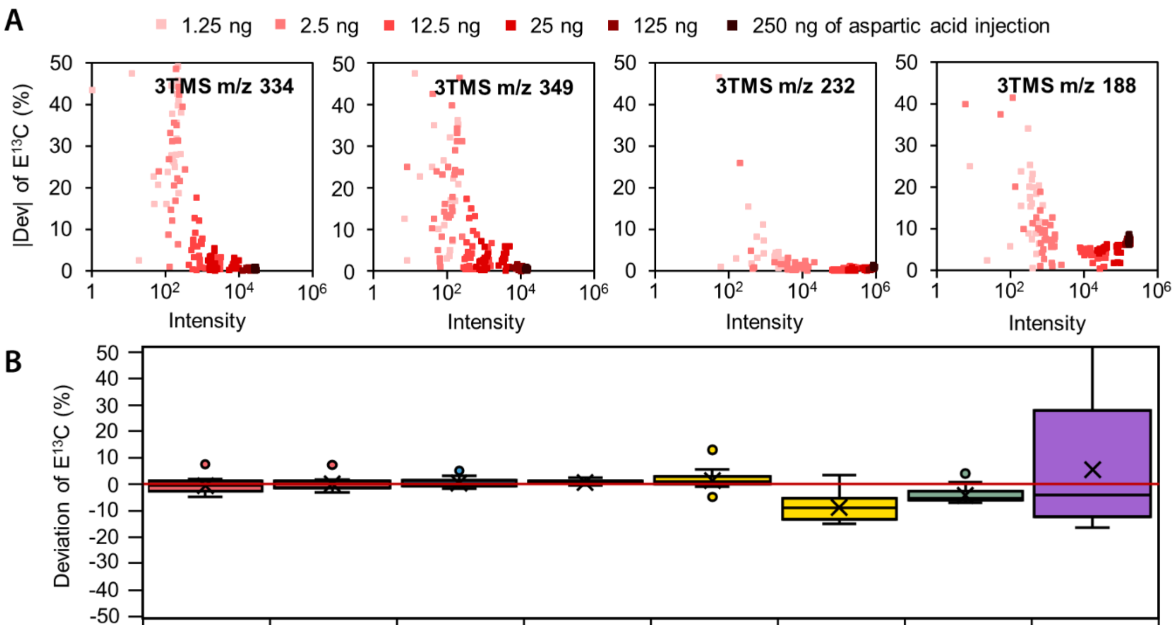


| m/z | 349 | 334 | 306 | 232 | 218 | 202 | 188 | 205 |
|---|---|---|---|---|---|---|---|---|
| Carbons | 1,2,3,4 | 1,2,3,4 | 2,3,4 | 2,3,4 | 1,2 | 1,2 | 2,3 | 3,4 |
| Mean (%) | -1.12 | -0.50 | -0.12 | 0.07 | 0.82 | -9.31 | -4.72 | 4.79 |
| SD (%) | 2.39 | 1.76 | 1.38 | 0.56 | 2.67 | 4.22 | 2.53 | -21.76 |

**Figure S 7. Accuracy and precision of $E^{13}C$ determination of aspartic acid (3TMS) by GC-EI-MS.** Equal mixtures of four positional $^{13}C$-labeled aspartic acid standards in different isotopic dilutions by natural aspartic acid and at different concentrations (Table S2) were measured by GC-EI-MS. Deviations of $E^{13}C$, i.e. measured $E^{13}C$ subtracted from expected $E^{13}C$, of the specified fragment ions were analyzed. (A) $E^{13}C$ absolute value deviations of selected fragments depended on fragment specific abundances, i.e. the sum of all isotopologue abundances, and amounts of injected aspartic acid. Most fragments provided accurate enrichment information with injections > 125 ng aspartic acid. Fragment m/z 232 $E^{13}C$ has high accuracy for injections higher 12.5 ng. Fragment m/z 188 shows absolute deviation of $E^{13}C$ of about 5% also for high fragment intensity and injections up to 250 ng. (B) Box-plot representation with means, and standard deviations (SD) of $E^{13}C$ deviations from selected fragments defined by nominal mass to charge ratio (m/z). 36 different mixtures were analyzed by 4 technical replicates injecting 25 ng aspartic acid. Indicated mean deviations and SD were determined as parameters of accuracy and precision, respectively. All fragments were analyzed by GC-MS measurements in split-less mode.

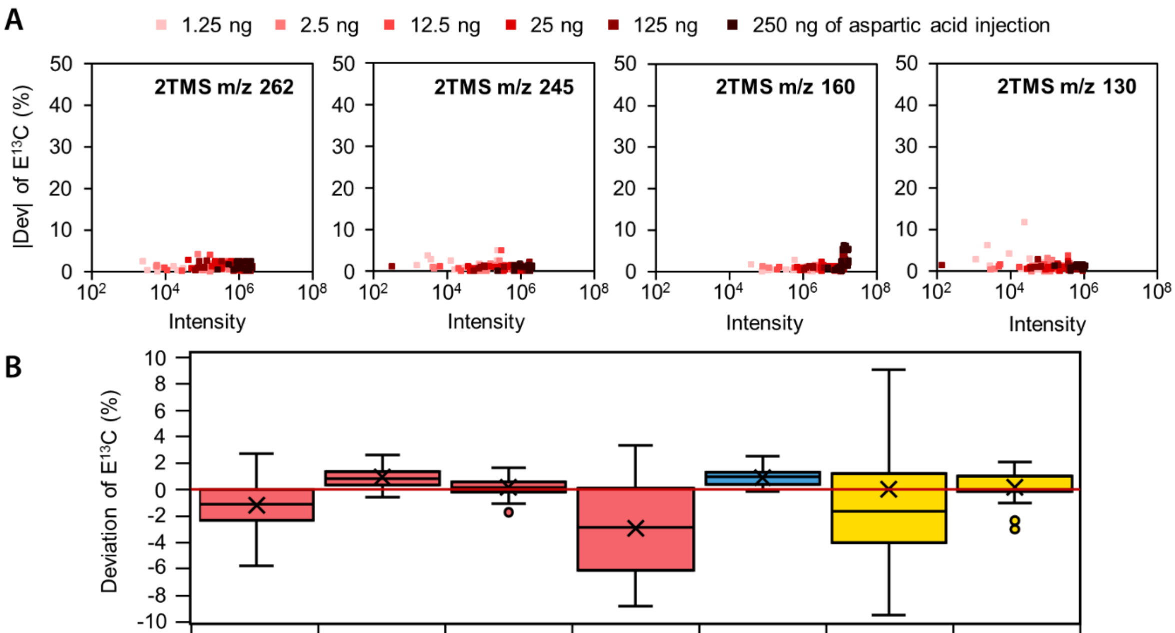


| m/z | 278 | 262 | 245 | 188 | 160 | 202 | 130 |
|---|---|---|---|---|---|---|---|
| Carbons | 1,2,3,4 | 1,2,3,4 | 1,2,3,4 | 1,2,3,4 | 2,3,4 | 1,2 | 1,2 |
| Mean (%) | -1.14 | 0.87 | 0.14 | -2.78 | 0.83 | 0.04 | 0.13 |
| SD (%) | 1.63 | 0.75 | 0.69 | 2.95 | 0.62 | 9.53 | 1.12 |

**Figure S 8. Accuracy of $E^{13}C$ determination of aspartic acid (2TMS) by GC-APCI-MS.** Equal mixtures of four positional $^{13}C$-labeled aspartic acid standards in different isotopic dilutions by natural aspartic acid and at different concentrations (Table S2) were measured by GC-APCI-MS. Deviations of $E^{13}C$, i.e. measured $E^{13}C$ subtracted from expected $E^{13}C$, of the specified fragment ions were analyzed. (A) $E^{13}C$ absolute value deviations of selected fragments depended on fragment specific abundances, i.e. the sum of all isotopologue abundances, and amounts of injected aspartic acid. For fragments m/z 262, m/z 245 and m/z 130, accuracy of $E^{13}C$ was mostly independent from fragment intensity. Fragment m/z 160 showed increased absolute value deviation of $E^{13}C$ for injections higher 125 ng due to saturation. (B) Box-plot representation with means, and standard deviations (SD) of $E^{13}C$ deviations from selected fragments defined by nominal mass to charge ratio (m/z). 36 different mixtures were analyzed by 4 technical replicates injecting 25 ng aspartic acid. Indicated mean deviations and SD were determined as parameters of accuracy and precision, respectively. All fragments were analyzed by GC-MS measurements in split-less mode.

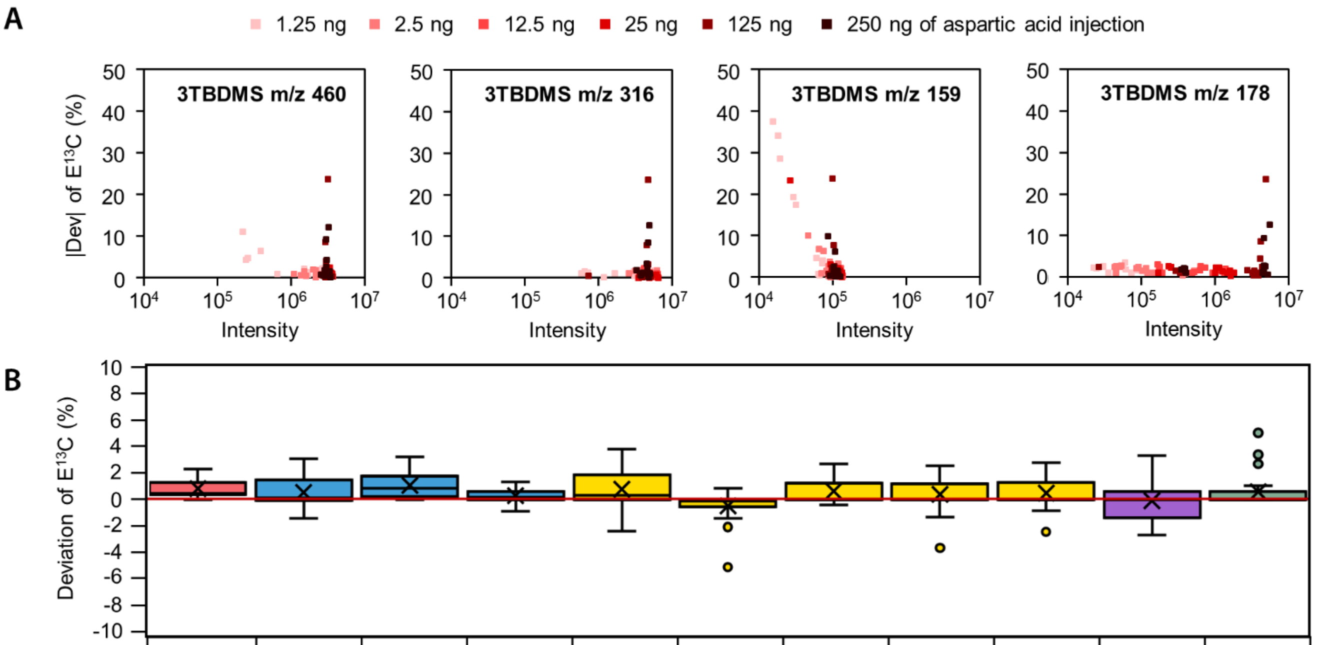


| m/z | 460 | 432 | 390 | 316 | 376 | 302 | 286 | 244 | 188 | 159 | 178 |
|---|---|---|---|---|---|---|---|---|---|---|---|
| Carbons | 1,2,3,4 | 2,3,4 | 2,3,4 | 2,3,4 | 1,2 | 1,2 | 1,2 | 1,2 | 1,2 | 3,4 | 2 |
| Mean (%) | 0.81 | 0.52 | 1.02 | 0.29 | 0.73 | -0.40 | 0.60 | 0.37 | 0.47 | -0.06 | 0.57 |
| SD (%) | 0.62 | 0.94 | 0.94 | 0.46 | 1.25 | 0.95 | 0.93 | 1.06 | 1.07 | 1.36 | 1.17 |

**Figure S 9. Accuracy of $E^{13}C$ determination of aspartic acid (3TBDMS) by GC-APCI-MS.** Equal mixtures of four positional $^{13}C$-labeled aspartic acid standards in different isotopic dilutions by natural aspartic acid and at different concentrations (Table S2) were measured by GC-APCI-MS. Deviations of $E^{13}C$, i.e. measured $E^{13}C$ subtracted from expected $E^{13}C$, of the specified fragment ions were analyzed. (A) $E^{13}C$ absolute value deviations of selected fragments depended on fragment specific abundances, i.e. the sum of all isotopologue abundances, and amounts of injected aspartic acid. Most fragments provided accurate enrichment information with injections > 12.5 ng aspartic acid. Saturation is reached for injections higher 125 ng leading to inaccuracy in $E^{13}C$ determination. (B) Box-plot representation with means, and standard deviations (SD) of $E^{13}C$ deviations from selected fragments defined by nominal mass to charge ratio (m/z). 36 different mixtures were analyzed by 4 technical replicates injecting 25 ng aspartic acid. Indicated mean deviations and SD were determined as parameters of accuracy and precision, respectively. All fragments were analyzed by GC-MS measurements in split-less mode.

**Table S 2. Fragment ion validation of trimethylsilylated and tert-butyldimethylsilylated derivatives of aspartic acid.** Aspartic acid was subjected to either trimethylsilylation or tert-butyldimethylsilylation followed by analysis with GC-EI-MS and GC-APCI-MS. Detection of aspartic acid derivatives 3TMS, 2TMS and 3TBDMS was possible using GC-EI-MS. Additionally to these, aspartic acid derivative 2TBDMS was detected by GC-APCI-MS. Included carbon atoms and number of potentially *in vivo* labeled $^{13}$C atoms were defined through analysis of positional labeled standards, natural, [U-$^{13}$C], [1-$^{13}$C], [2-$^{13}$C], [3-$^{13}$C], and [4-$^{13}$C]aspartic acids. Molecular formula and exact mass were predicted through *in silico* fragmentation analyses. Predicted exact mass was compared with measured exact mass from GC-APCI-MS (mean ± standard deviation (SD) from 3 individual experiments). Detected mass accuracy was higher for most fragments than the previously published[1] mean deviation of 1.3 mDa ± 0.8 mDa. Mean deviation of E$^{13}$C in GC-EI-MS and GC-APCI-MS was determined using mixtures of positional labeled aspartic acid standards in different ratios and at isotopic concentrations adjusted by natural aspartic acid (25 ng of aspartic acid injection, 36 different mixtures, at least 3 technical replicates, ND = not detected).

| Analyte | Fragment (m/z) | Included carbons | Number of *in vivo* labelled $^{13}$C atoms | Molecular formula (predicted) | Exact mass (predicted, Da) | Exact mass (measured, Da) | Mass accuracy (mDa) | Mean deviation of E$^{13}$C by GC-EI-MS ± SD (%) | Mean deviation of E$^{13}$C by GC-APCI-MS ± SD (%) |
|---|---|---|---|---|---|---|---|---|---|
| 3TMS | 350 | 1,2,3,4 | 4 | $C_{13}H_{32}N_1O_4Si_3$ | 350.1634 | 350.1630 | 0.4 | ND | 1.1 ± 0.6 |
| 3TMS | 349 | 1,2,3,4 | 4 | $C_{13}H_{31}N_1O_4Si_3$ | 349.1555 | 349.1550 | 0.5 | -1.1 ± 2.4 | ND |
| 3TMS | 334 | 1,2,3,4 | 4 | $C_{12}H_{28}N_1O_4Si_3$ | 334.1321 | 334.1322 | -0.1 | -0.5 ± 1.8 | 0.2 ± 0.7 |
| 3TMS | 306 | 2,3,4 | 3 | $C_{11}H_{28}N_1O_3Si_3$ | 306.1372 | 306.1374 | -0.3 | -0.1 ± 1.4 | 0.0 ± 1.0 |
| 3TMS | 278 | 1,2,3,4 | 4 | $C_{10}H_{24}N_1O_4Si_2$ | 278.1238 | 278.1235 | 0.3 | ND | 0.6 ± 0.5 |
| 3TMS | 232 | 2,3,4 | 3 | $C_9H_{22}N_1O_2Si_2$ | 232.1184 | 232.1182 | 0.1 | 0.1 ± 0.6 | 0.7 ± 0.8 |
| 3TMS | 218 | 1,2 | 2 | $C_8H_{20}N_1O_2Si_2$ | 218.1027 | 218.1027 | 0.0 | 0.8 ± 2.7 | 0.4 ± 1.1 |
| 3TMS | 205 | 3,4 | 2 | $C_8H_{21}O_2Si_2$ | 205.1075 | 205.1073 | 0.2 | 4.8 ± 21.8 | 1.0 ± 1.0 |
| 3TMS | 202 | 1,2 | 2 | $C_7H_{16}N_1O_2Si_2$ | 202.0714 | 202.0716 | -0.2 | -9.3 ± 4.2 | 0.4 ± 0.9 |
| 3TMS | 188 | 2,3 | 2 | $C_8H_{22}N_1Si_2$ | 188.1285 | 188.1286 | -0.1 | -4.7 ± 2.5 | 0.4 ± 0.6 |
| 3TMS | 160 | 2,3,4 | 3 | $C_6H_{14}N_1O_2Si_1$ | 160.0788 | 160.0787 | 0.1 | ND | 0.9 ± 0.6 |
| 2TMS | 278 | 1,2,3,4 | 4 | $C_{10}H_{24}N_1O_4Si_2$ | 278.1238 | 278.1236 | 0.2 | ND | -1.1 ± 1.6 |
| 2TMS | 277 | 1,2,3,4 | 4 | $C_{10}H_{23}N_1O_4Si_2$ | 277.1160 | 277.1158 | 0.2 | ND | ND |
| 2TMS | 262 | 1,2,3,4 | 4 | $C_9H_{20}N_1O_4Si_2$ | 262.0925 | 262.0925 | 0.0 | 0.6 ± 6.2 | 0.9 ± 0.7 |
| 2TMS | 245 | 1,2,3,4 | 4 | $C_9H_{17}O_4Si_2$ | 245.0660 | 245.0656 | 0.4 | -10.6 ± 5.0 | 0.1 ± 0.7 |
| 2TMS | 202 | 1,2 | 2 | $C_7H_{16}N_1O_2Si_2$ | 202.0714 | 202.0719 | -0.4 | 0.6 ± 6.2 | 0.0 ± 9.6 |
| 2TMS | 188 | 1,2,3,4 | 4 | $C_7H_{14}N_1O_3Si_1$ | 188.0737 | 188.0736 | 0.1 | ND | 2.8 ± 3.0 |
| 2TMS | 160 | 2,3,4 | 3 | $C_6H_{14}N_1O_2Si_1$ | 160.0788 | 160.0787 | 0.1 | 0.6 ± 1.2 | 0.8 ± 0.6 |
| 2TMS | 130 | 1,2 | 2 | $C_4H_8N_1O_2Si_1$ | 130.0319 | 130.0316 | 0.3 | -2.1 ± 2.3 | 0.1 ± 1.1 |
| 3TBDMS | 476 | 1,2,3,4 | 4 | $C_{22}H_{50}N_1O_4Si_3$ | 476.3042 | 476.3034 | 0.8 | ND | -5.0 ± 2.3 |
| 3TBDMS | 460 | 1,2,3,4 | 4 | $C_{21}H_{46}N_1O_4Si_3$ | 460.2729 | 460.2721 | 0.8 | -0.5 ± 3.1 | 0.8 ± 0.6 |
| 3TBDMS | 432 | 2,3,4 | 3 | $C_{20}H_{46}N_1O_3Si_3$ | 432.2780 | 432.2775 | 0.5 | -7.8 ± 6.1 | 0.5 ± 1.0 |

| | | | | | | | | | |
|---|---|---|---|---|---|---|---|---|---|
| 3TBDMS | 418 | 1,2,3,4 | 4 | $C_{18}H_{40}N_1O_4Si_3$ | 418.2260 | 418.2254 | 0.6 | 0.4 ± 2.4 | -5.8 ± 2.7 |
| 3TBDMS | 390 | 2,3,4 | 3 | $C_{17}H_{40}N_1O_3Si_3$ | 390.2311 | 390.2303 | 0.7 | -4.2 ± 2.3 | 1.0 ± 0.9 |
| 3TBDMS | 376 | 1,2 | 2 | $C_{16}H_{38}N_1O_3Si_3$ | 376.2154 | 376.2151 | 0.3 | -6.3 ± 3.7 | 0.7 ± 1.3 |
| 3TBDMS | 316 | 2,3,4 | 3 | $C_{15}H_{34}N_1O_2Si_2$ | 316.2123 | 316.2117 | 0.6 | -0.7 ± 1.1 | 0.3 ± 0.5 |
| 3TBDMS | 302 | 1,2 | 2 | $C_{14}H_{32}N_1O_2Si_2$ | 302.1966 | 302.1961 | 0.5 | -0.5 ± 1.3 | -0.4 ± 1.0 |
| 3TBDMS | 287 | 1,2,3,4 | 4 | $C_{13}H_{29}N_1O_2Si_2$ | 287.1731 | ND | ND | 2.8 ± 7.4 | ND |
| 3TBDMS | 286 | 1,2 | 2 | $C_{13}H_{28}N_1O_2Si_2$ | 286.1653 | 286.1650 | 0.3 | ND | 0.6 ± 0.9 |
| 3TBDMS | 258 | 2,3,4 | 3 | $C_{11}H_{24}N_1O_2Si_2$ | 258.1340 | 258.1342 | -0.2 | -3.6 ± 3.0 | -1.4 ± 4.2 |
| 3TBDMS | 244 | 1,2 | 2 | $C_{10}H_{22}N_1O_2Si_2$ | 244.1184 | 244.1180 | 0.4 | -7.9 ± 3.9 | 0.4 ± 1.1 |
| 3TBDMS | 202 | 1,2 | 2 | $C_9H_{20}N_1O_2Si_1$ | 202.1258 | 202.1258 | 0.0 | -16.6 ± 11.9 | -3.8 ± 25.2 |
| 3TBDMS | 188 | 1,2 | 2 | $C_6H_{14}N_1O_2Si_2$ | 188.0558 | 188.0556 | 0.2 | ND | 0.5 ± 1.1 |
| 3TBDMS | 178 | 2 | 1 | $C_5H_{16}N_1O_2Si_2$ | 178.0714 | 178.0713 | 0.1 | ND | 0.6 ± 1.2 |
| 3TBDMS | 159 | 3,4 | 2 | $C_7H_{15}O_2Si_1$ | 159.0836 | 159.0835 | 0.1 | ND | -0.1 ± 1.4 |
| 3TBDMS | 117 | 3,4 | 2 | $C_4H_9O_2Si_1$ | 117.0366 | 117.0365 | 0.2 | -3.5 ± 8.0 | -4.6 ± 4.1 |
| 2TBDMS | 476 | 1,2,3,4 | 4 | $C_{22}H_{50}N_1O_4Si_3$ | 476.3042 | 476.3042 | 0.0 | ND | -2.4 ± 2.7 |
| 2TBDMS | 460 | 1,2,3,4 | 4 | $C_{21}H_{46}N_1O_4Si_3$ | 460.2729 | 460.2720 | 0.9 | ND | 0.8 ± 0.7 |
| 2TBDMS | 418 | 1,2,3,4 | 4 | $C_{18}H_{40}N_1O_4Si_3$ | 418.2260 | 418.2254 | 0.6 | ND | 0.7 ± 0.6 |
| 2TBDMS | 408 | 1,2,3,4 | 4 | $C_{16}H_{42}N_1O_3Si_4$ | 408.2236 | 408.2226 | 1.0 | ND | 0.4 ± 1.0 |
| 2TBDMS | 362 | 1,2,3,4 | 4 | $C_{16}H_{36}N_1O_4Si_2$ | 362.2177 | 362.2178 | 0.0 | ND | -4.4 ± 2.6 |
| 2TBDMS | 346 | 1,2,3,4 | 4 | $C_{15}H_{32}N_1O_4Si_2$ | 346.1864 | 346.1861 | 0.4 | ND | 1.9 ± 1.6 |
| 2TBDMS | 316 | 2,3,4 | 3 | $C_{15}H_{34}N_1O_2Si_2$ | 316.2123 | 316.2146 | -2.4 | ND | 2.4 ± 3.3 |
| 2TBDMS | 304 | 1,2,3,4 | 4 | $C_{12}H_{26}N_1O_4Si_2$ | 304.1395 | 304.1390 | 0.5 | ND | 0.8 ± 0.8 |
| 2TBDMS | 276 | 2,3,4 | 3 | $C_{11}H_{26}N_1O_3Si_2$ | 276.1446 | 276.1442 | 0.4 | ND | 0.6 ± 0.6 |
| 2TBDMS | 262 | 1,2 | 2 | $C_{10}H_{24}N_1O_3Si_2$ | 262.1289 | 262.1286 | 0.3 | ND | -0.6 ± 3.2 |
| 2TBDMS | 216 | 2 | 1 | $C_9H_{22}N_1O_1Si_2$ | 216.1234 | 216.1236 | -0.1 | ND | 0.7 ± 2.3 |
| 2TBDMS | 202 | 2,3,4 | 3 | $C_9H_{20}N_1O_2Si_1$ | 202.1258 | 202.1254 | 0.3 | ND | 0.6 ± 0.5 |
| 2TBDMS | 188 | 1,2 | 2 | $C_8H_{18}N_1O_2Si_1$ | 188.1101 | 188.1099 | 0.3 | ND | -3.1 ± 2.9 |
| 2TBDMS | 162 | 2,3,4 | 3 | $C_5H_{12}N_1O_3Si_1$ | 162.0581 | 162.0578 | 0.3 | ND | 0.5 ± 0.8 |
| 2TBDMS | 158 | 2,3 | 2 | $C_8H_{20}N_1Si_1$ | 158.1360 | 158.1357 | 0.3 | ND | 0.4 ± 0.6 |
| 2TBDMS | 130 | 1,2 | 2 | $C_4H_8N_1O_2Si_1$ | 130.0319 | 130.0316 | 0.3 | ND | 0.3 ± 1.0 |
| 2TBDMS | 100 | 2,3 | 2 | $C_4H_{10}N_1Si_1$ | 100.0577 | 100.0575 | 0.2 | ND | 0.4 ± 0.6 |

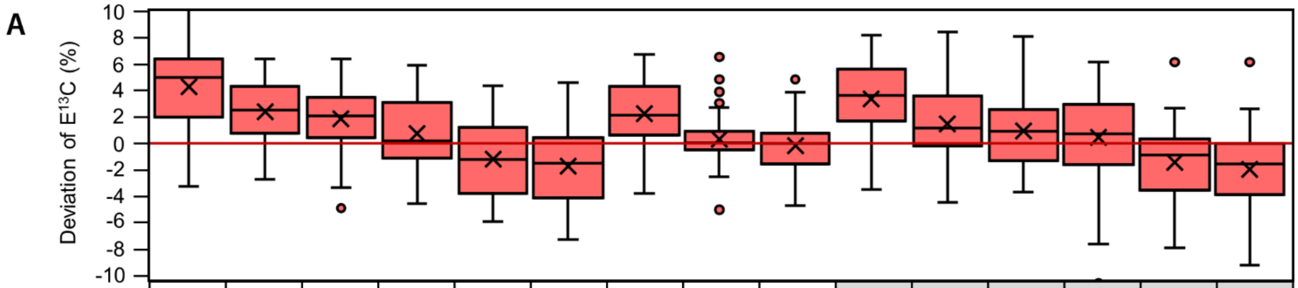


| m/z 1,2,3,4 | 350 | 350 | 350 | 334 | 334 | 334 | 278 | 278 | 278 | 262 | 262 | 262 | 245 | 245 | 245 |
|---|---|---|---|---|---|---|---|---|---|---|---|---|---|---|---|
| m/z 2,3,4 | 306 | 232 | 160 | 306 | 232 | 160 | 306 | 232 | 160 | 306 | 232 | 160 | 306 | 232 | 160 |
| **Position** | **1** | **1** | **1** | **1** | **1** | **1** | **1** | **1** | **1** | **1** | **1** | **1** | **1** | **1** | **1** |
| Mean (%) | 4.36 | 2.43 | 1.90 | 0.79 | -1.14 | -1.67 | 2.31 | 0.38 | -0.15 | 3.44 | 1.51 | 098 | 0.52 | -1.42 | -1.94 |
| SD (%) | 3.36 | 2.22 | 2.52 | 2.81 | 3.19 | 3.12 | 2.53 | 2.22 | 2.04 | 3.05 | 2.97 | 2.76 | 3.65 | 2.9 | 3.02 |

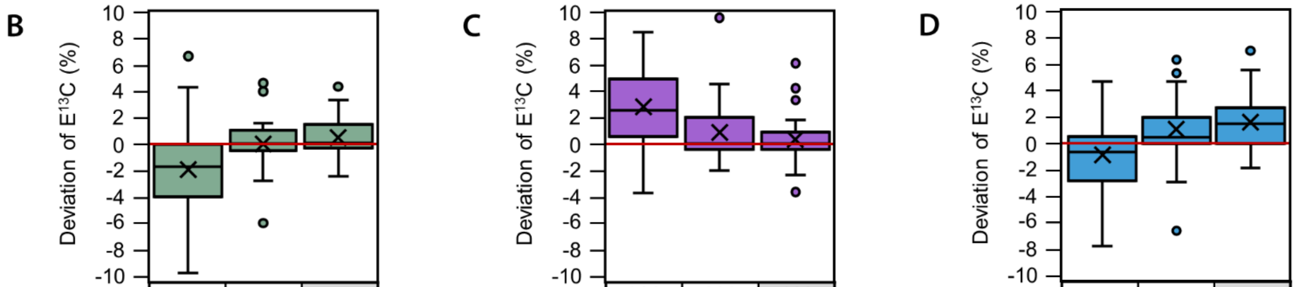


| m/z 2,3,4 | 306 | 232 | 160 |
|---|---|---|---|
| m/z 3,4 | 205 | 205 | 205 |
| **Position** | **2** | **2** | **2** |
| Mean (%) | -1.67 | 0.03 | 0.64 |
| SD (%) | 3.21 | 1.94 | 1.57 |

| m/z 2,3,4 | 306 | 232 | 160 |
|---|---|---|---|
| m/z 3,4 | 205 | 205 | 205 |
| m/z 2,3 | 188 | 188 | 188 |
| **Position** | **3** | **3** | **3** |
| Mean (%) | 2.87 | 0.93 | 0.40 |
| SD (%) | 3.10 | 2.15 | 1.83 |

| m/z 2,3,4 | 306 | 232 | 160 |
|---|---|---|---|
| m/z 2,3 | 188 | 188 | 188 |
| **Position** | **4** | **4** | **4** |
| Mean (%) | -0.95 | 0.75 | 1.36 |
| SD (%) | 2.84 | 2.40 | 1.88 |

**Figure S 10. Possibilities to calculate all C-positional E$^{13}$C of aspartic acid using derivatives 3TMS and 2TMS by GC-APCI-MS.** Accuracy and precision of positional E$^{13}$C from aspartic acid 3TMS determined by GC-APCI-MS. Positional E$^{13}$C was calculated applying equations **Error! Reference source not found.**), **Error! Reference source not found.**), **Error! Reference source not found.**) and **Error! Reference source not found.**) to E$^{13}$C of the indicated fragment ions (m/z, grey shaded fragments are originating from aspartic acid 2TMS, white underlaid from aspartic acid 3TMS). Distribution analyses of E$^{13}$C deviations, i.e. calculated E$^{13}$C subtracted from expected, is displayed across 36 different mixtures with 4 technical replicates (cf. Legend to Figure 2). Amount of total aspartic acid injected equaled 25 ng. E$^{13}$C of fragment ion m/z 232 was determined by split measurements at split ratio 1:5. E$^{13}$Cs of all other fragment ions were determined by analyses in split-less mode.

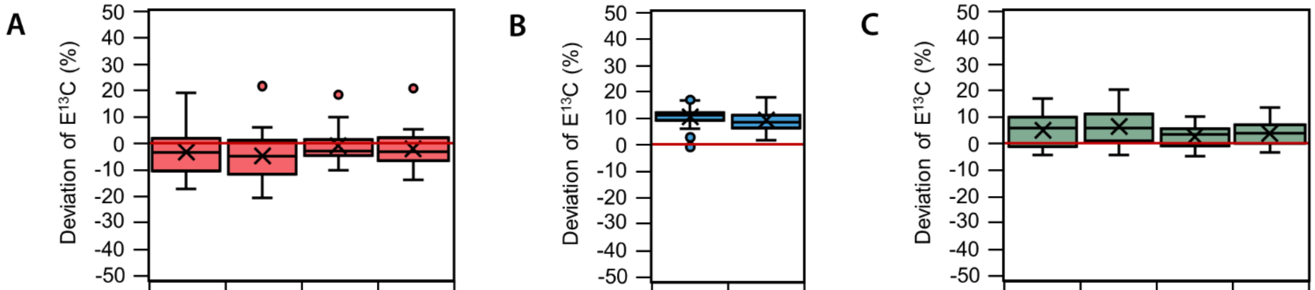


**A**

| | | | | |
|---|---|---|---|---|
| m/z 1,2,3,4 | 349 | 349 | 334 | 334 |
| m/z 2,3,4 | 306 | 232 | 306 | 232 |
| **Position** | **1** | **1** | **1** | **1** |
| Mean (%) | -3.42 | -4.70 | -0.92 | -2.20 |
| SD (%) | 8.03 | 8.42 | 6.20 | 6.13 |

**B**

| | | |
|---|---|---|
| m/z 2,3,4 | 306 | 232 |
| m/z 2,3 | 188 | 188 |
| **Position** | **4** | **4** |
| Mean (%) | -3.42 | -4.70 |
| SD (%) | 8.03 | 8.42 |

**C**

| | | | | |
|---|---|---|---|---|
| m/z 1,2 | 218 | 218 | 218 | 218 |
| 1-C | 349/306 | 349/232 | 334/306 | 334/232 |
| **Position** | **2** | **2** | **2** | **2** |
| Mean (%) | 4.84 | 6.11 | 2.34 | 3.61 |
| SD (%) | 6.33 | 6.79 | 4.25 | 4.09 |

**Figure S 11. Options of positional $E^{13}C$ calculations of aspartic acid using GC-EI-MS.** Accuracy and precision of positional $E^{13}C$ from aspartic acid 3TMS determined by GC-EI-MS. Positional $E^{13}C$ for 1-C and 4-C was calculated applying equations **Error! Reference source not found.**) and **Error! Reference source not found.**) to $E^{13}C$ of the indicated fragment ions (m/z). $E^{13}C$ of 2-C (C) was determined by subtracting $E^{13}C$ of 1-C, calculated based on fragment ions with indicated m/z, from $E^{13}C$ of fragment m/z 218 including 1,2-C. Distribution analyses of $E^{13}C$ deviations, i.e. calculated $E^{13}C$ subtracted from expected, are displayed across 36 different mixtures with 4 technical replicates (cf. Legend to Figure 2). Amount of total aspartic acid injected equaled 25 ng. $E^{13}Cs$ of all fragments were determined by analyses in split-less mode. Positional $E^{13}C$ of 4-C is underestimated due to mass isotopologue distribution interference of $E^{13}C$ of m/z 188.

A

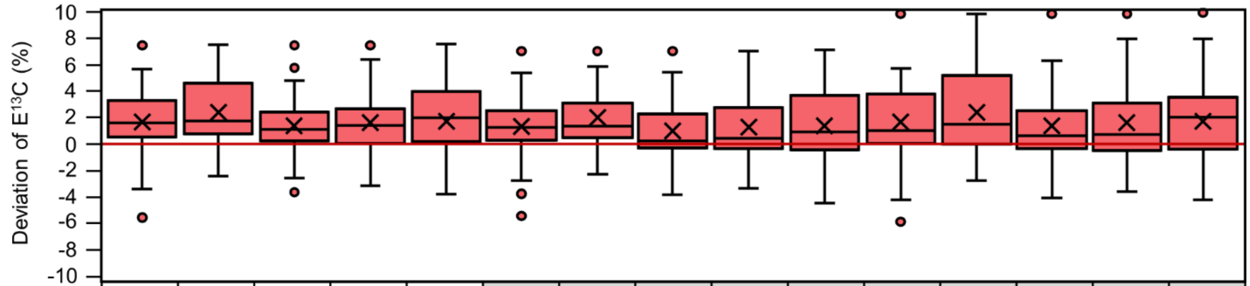


| | | | | | | | | | | | | | | | |
|---|---|---|---|---|---|---|---|---|---|---|---|---|---|---|---|
| m/z 1,2,3,4 | 460 | 460 | 460 | 460 | 460 | 418 | 418 | 418 | 418 | 418 | 304 | 304 | 304 | 304 | 304 |
| m/z 2,3,4 | 432 | 316 | 276 | 202 | 162 | 432 | 316 | 276 | 202 | 162 | 432 | 316 | 276 | 202 | 162 |
| **Position** | **1** | **1** | **1** | **1** | **1** | **1** | **1** | **1** | **1** | **1** | **1** | **1** | **1** | **1** | **1** |
| Mean (%) | 1.66 | 2.36 | 1.35 | 1.61 | 1.70 | 1.30 | 2.00 | 0.99 | 1.25 | 1.34 | 1.66 | 2.36 | 1.34 | 1.60 | 1.70 |
| SD (%) | 2.50 | 2.50 | 2.27 | 2.48 | 3.16 | 2.35 | 2.39 | 2.20 | 2.40 | 3.12 | 2.93 | 3.21 | 2.78 | 3.12 | 3.64 |

B

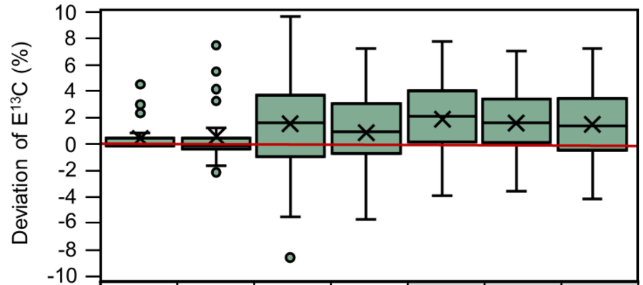


| | | | | | | | |
|---|---|---|---|---|---|---|---|
| m/z 2,3,4 | - | - | 432 | 316 | 276 | 202 | 162 |
| m/z 3,4 | - | - | 159 | 159 | 159 | 159 | 159 |
| m/z 2 | 178 | 216 | - | - | - | - | - |
| **Position** | **2** | **2** | **2** | **2** | **2** | **2** | **2** |
| Mean (%) | 0.57 | 0.68 | 1.70 | 0.99 | 2.01 | 1.75 | 1.66 |
| SD (%) | 1.17 | 2.30 | 4.05 | 3.00 | 2.74 | 2.50 | 2.72 |

C

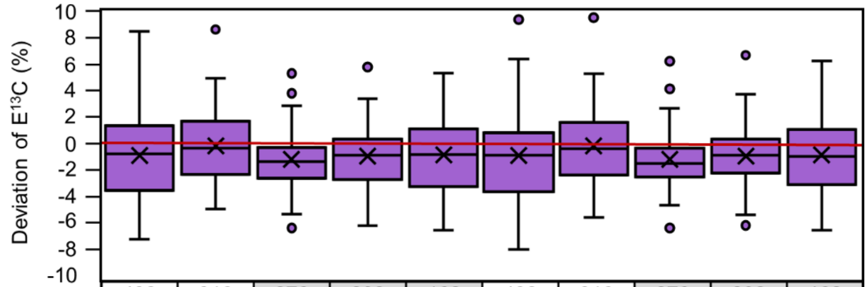


| | | | | | | | | | | |
|---|---|---|---|---|---|---|---|---|---|---|
| m/z 2,3,4 | 432 | 316 | 276 | 202 | 162 | 432 | 316 | 276 | 202 | 162 |
| m/z 2,3 | 158 | 158 | 158 | 158 | 158 | 100 | 100 | 100 | 100 | 100 |
| m/z 3,4 | 159 | 159 | 159 | 159 | 159 | 159 | 159 | 159 | 159 | 159 |
| **Position** | **3** | **3** | **3** | **3** | **3** | **3** | **3** | **3** | **3** | **3** |
| Mean (%) | -0.87 | -0.16 | -1.18 | -0.92 | -0.83 | -0.86 | -0.16 | -1.17 | -0.91 | -0.82 |
| SD (%) | 3.86 | 3.05 | 2.47 | 2.47 | 2.73 | 3.98 | 3.15 | 2.60 | 2.56 | 2.82 |

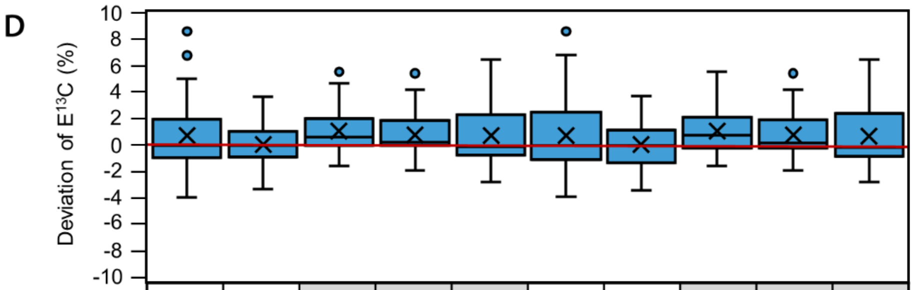


| | | | | | | | | | | |
|---|---|---|---|---|---|---|---|---|---|---|
| m/z 2,3,4 | 432 | 316 | 276 | 202 | 162 | 432 | 316 | 276 | 202 | 162 |
| m/z 2,3 | 158 | 158 | 158 | 158 | 158 | 100 | 100 | 100 | 100 | 100 |
| **Position** | **4** | **4** | **4** | **4** | **4** | **4** | **4** | **4** | **4** | **4** |
| Mean (%) | 0.74 | 0.04 | 1.05 | 0.80 | 0.70 | 0.74 | 0.03 | 1.05 | 0.79 | 0.70 |
| SD (%) | 2.67 | 1.71 | 1.62 | 1.61 | 2.53 | 2.76 | 1.77 | 1.69 | 1.62 | 2.55 |

**Figure S 12. Options of positional $E^{13}C$ determinations of aspartic acid by GC-APCI-MS after TBDMS derivatization.** Accuracy and precision of positional $E^{13}C$ from aspartic acid 2TBDMS and 3TBDMS determined by GC-APCI-MS. Positional $E^{13}C$ was calculated with formulas 1, 2, 3 and 4 for 1-C (A), 2-C (B), 3-C (C) and 4-C (D), respectively, using the indicated fragments (grey shaded fragments originate from aspartic acid 2TBDMS, white underlaid fragments are from aspartic acid 3TBDMS). For 2-C (B), fragments that only included 2-C are displayed (m/z 178, m/z 216). Distributions of $E^{13}C$ deviation of positional $E^{13}Cs$, i.e. calculated $E^{13}C$ subtracted from expected $E^{13}C$, are displayed for 36 different mixtures with 4 technical replicates (cf. Legend to Figure 2). The injected amount of total aspartic acid equaled 25 ng. Indicated mean deviation and standard deviation (SD) were determined as parameters for accuracy and precision, respectively.

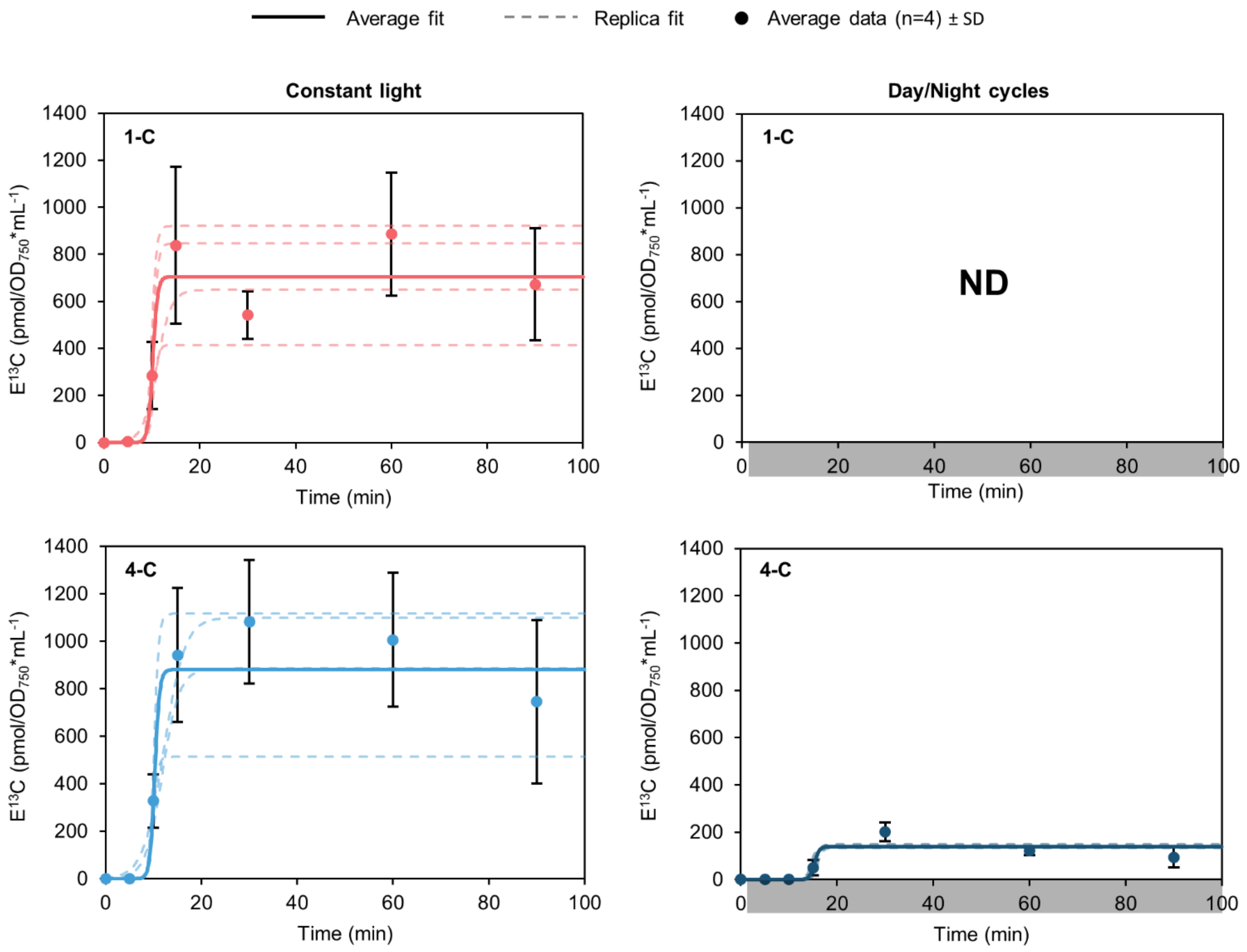


**Figure S 13. Sigmoidal curve fitting of aspartate 1-C and 4-C labeling within *Synechocystis* cultures during the day and the night.** *Synechocystis* cells were cultivated photoautotrophically with 5% $CO_2$-enriched air. Cells were either cultivated in constant light (CL) or a 12 h:12 h day/night photoperiod (DN). For DN, the $^{13}CO_2$ labelling pulse was applied after transition to the night. Samples were taken 5 to 90 min after the labelling pulse. Samples were analyzed by GC-MS. Data points represent average of 4 replicates ± SD. Curve fitting was done for $E^{13}C$ of aspartate 1-C and 4-C to characterize contributions of RUBISCO and PEPC activities, respectively. Sigmoidal fits are displayed for the average data points (solid line) and the single replicates (n=4, dashed lines). No $E^{13}C$ was detected in aspartate 1-C during the night.


1. Strehmel, N.; Kopka, J.; Scheel, D.; Böttcher, C., Annotating unknown components from GC/EI-MS-based metabolite profiling experiments using GC/APCI(+)-QTOFMS. *Metabolomics* **2014,** *10* (2), 324-336.
